# Dachsous and Fat coordinately repress the Dachs-Dlish-Approximated complex to control growth

**DOI:** 10.1101/2024.06.18.599638

**Authors:** Hitoshi Matakatsu, Richard G. Fehon

## Abstract

Two protocadherins, Dachsous (Ds) and Fat (Ft), regulate organ growth *in Drosophila* via the Hippo pathway. Ds and Ft bind heterotypically to regulate the abundance and subcellular localization of a ‘core complex’ consisting of Dachs, Dlish and Approximated. This complex localizes to the junctional cortex where it promotes growth by repressing the pathway kinase Warts. Ds is believed to promote growth by recruiting and stabilizing the core complex at the junctional cortex, while Ft represses growth by promoting degradation of core complex components.

Here, we examine the functions of intracellular domains of Ds and Ft and their relationship to the core complex. While Ds promotes accumulation of the core complex proteins in cortical puncta, it is not required for core complex assembly. Indeed, the core complex assembles maximally in the absence of both Ds and Ft. Furthermore, while Ds promotes growth in the presence of Ft, it represses growth in the absence of Ft by removing the core complex from the junctional cortex. Ft similarly recruits core complex components, however it normally promotes their degradation. Our findings reveal that Ds and Ft constrain tissue growth by repressing the default ’on’ state of the core complex.

## Introduction

The ability to precisely control the subcellular localization of proteins, particularly at the cell cortex, is critical to the function of diverse intra- and intercellular signaling pathways. This property is well demonstrated by the Hippo pathway, which controls organ growth during development (Zheng and Pan, 2019; Karaman and Halder, 2018; Boggiano and Fehon, 2012). Hippo pathway kinases, including Tao, Hippo and Warts (Wts), function in a cascade to promote phosphorylation of the transcriptional co-factor Yorkie (Yki), resulting in Yki repression and decreased transcription of its pathway targets. While the kinases are normally diffusely cytoplasmic, previous studies have revealed that their assembly and activity are controlled by cortically localized protein complexes (Boggiano and Fehon, 2012; Karaman and Halder, 2018). Expanded, a FERM-domain protein, associates with the transmembrane protein Crumbs at the junctional cortex to assemble and activate the core kinases (Ling et al., 2010; Chen et al., 2010; Robinson et al., 2010). In parallel, Merlin (another FERM domain protein) and Kibra together assemble the same kinases at the apical medial cortex (Su et al., 2017).

Perhaps the most enigmatic cortically localized protein complex controlling Hippo pathway output is the Dachsous (Ds)-Fat (Ft) system. Ds and Ft are giant protocadherins that regulate planar cell polarity, proximo-distal patterning and organ growth (Mahoney et al., 1991; Clark et al., 1995; Bryant et al., 1988; Adler et al., 1998; reviewed in Blair and McNeill, 2018; Strutt and Strutt, 2021). Ft and Ds bind extracellularly via heterotypic interaction (Matakatsu and Blair, 2004; Ma et al., 2003; Strutt and Strutt, 2002). Opposing proximal-distal expression gradients of Ds and Four-jointed (Fj), a Golgi resident kinase that modulates Ds-Ft binding (Ma et al., 2003; Zeidler et al., 2000; Brittle et al., 2010; Simon et al., 2010), result in a planar polarized subcellular distribution of Ds on distal junctions and Ft on proximal junctions in wing discs (Brittle et al., 2012; Bosveld et al., 2012; Ambegaonkar et al., 2012).

Ft and Ds are thought to control tissue growth primarily via an atypical myosin, Dachs at the junctional cortex (Mao et al., 2006; Rogulja et al., 2008). Dachs promotes growth by inhibiting Wts function, by degrading Wts and/or regulating Wts conformational state (Cho et al., 2006; Vrabioiu and Struhl, 2015). Two additional proteins, <u>D</u>achs <u>Li</u>gand with <u>SH</u>3 domains (Dlish, aka Vamana) and a DHHC palmitoyltransferase Approximated (App) are required for Dachs function and localization to the junctional cortex (Matakatsu and Blair, 2008; Matakatsu et al., 2017; Zhang et al., 2016; Misra and Irvine, 2016). Dlish forms a complex with Dachs, Ft and Ds when expressed in S2 cells, and Dlish and Dachs are interdependent for cortical localization *in vivo* (Zhang et al., 2016; Misra and Irvine, 2016). App palmitoylates both Dlish and Ft (Matakatsu et al., 2017; Zhang et al., 2016). Palmitoylation often increases the affinity of a protein for the plasma membrane (Linder and Deschenes, 2007), leading to the suggestion that App promotes Dlish-dependent Dachs localization to the junctional cortex (Zhang et al., 2016). Since the accumulation of Dachs at the cortex is influenced by Ds-Ft planar polarity, it has been proposed that Ds recruits Dachs to the distal edge to promote growth while Ft promotes Dachs degradation at the proximal edge to suppress growth (Rogulja et al., 2008; Bosveld et al., 2012; Brittle et al., 2012). However, this model seems inconsistent with the observation that *ds ft* double mutants have a much stronger overgrowth phenotype than either single mutant alone, which instead suggests that Ft and Ds work synergistically to repress growth (Matakatsu and Blair, 2006).

To understand how Ds-Ft signaling regulates the abundance, localization and function of Dachs, Dlish and App (hereafter called the core complex) at the junctional cortex, we explored the functions of the intracellular domains of Ds and Ft. We find that members of the core complex, in particular App, depend on the Ds ICD to accumulate in cortical puncta and promote tissue growth. In contrast, in the absence of its ligand Ft, Ds (via its ICD) promotes mislocalization and inactivation of the core complex. Strikingly, loss of both Ds and Ft allows the core complex to accumulate to very high levels at the junctional cortex, indicating that neither Ds nor Ft is required for its cortical localization. Instead, our results suggest that Ds and Ft organize core complex proteins within the junctional cortex into discrete puncta with opposing consequences: Ft promotes degradation of the core complex whereas Ds protects it from degradation. Together these findings reveal previously unrecognized mechanisms by which Ds and Ft control the subcellular localization and abundance of Dachs, Dlish and App at the junctional cortex and thereby constrain tissue growth.

## Results

### Ft and Ds act synergistically to repress Yki activity

Current models propose that Ds and Ft function antagonistically. However, mutation of either gene causes overgrowth of wing discs (Fig. 1A, B and F) and combining mutations in both genes produces much greater overgrowth (Fig. 1G, Bryant et al., 1988; Matakatsu and Blair, 2006). Together, these results suggest that rather than functioning antagonistically Ds and Ft function synergistically to repress tissue growth.

**Figure 1.**
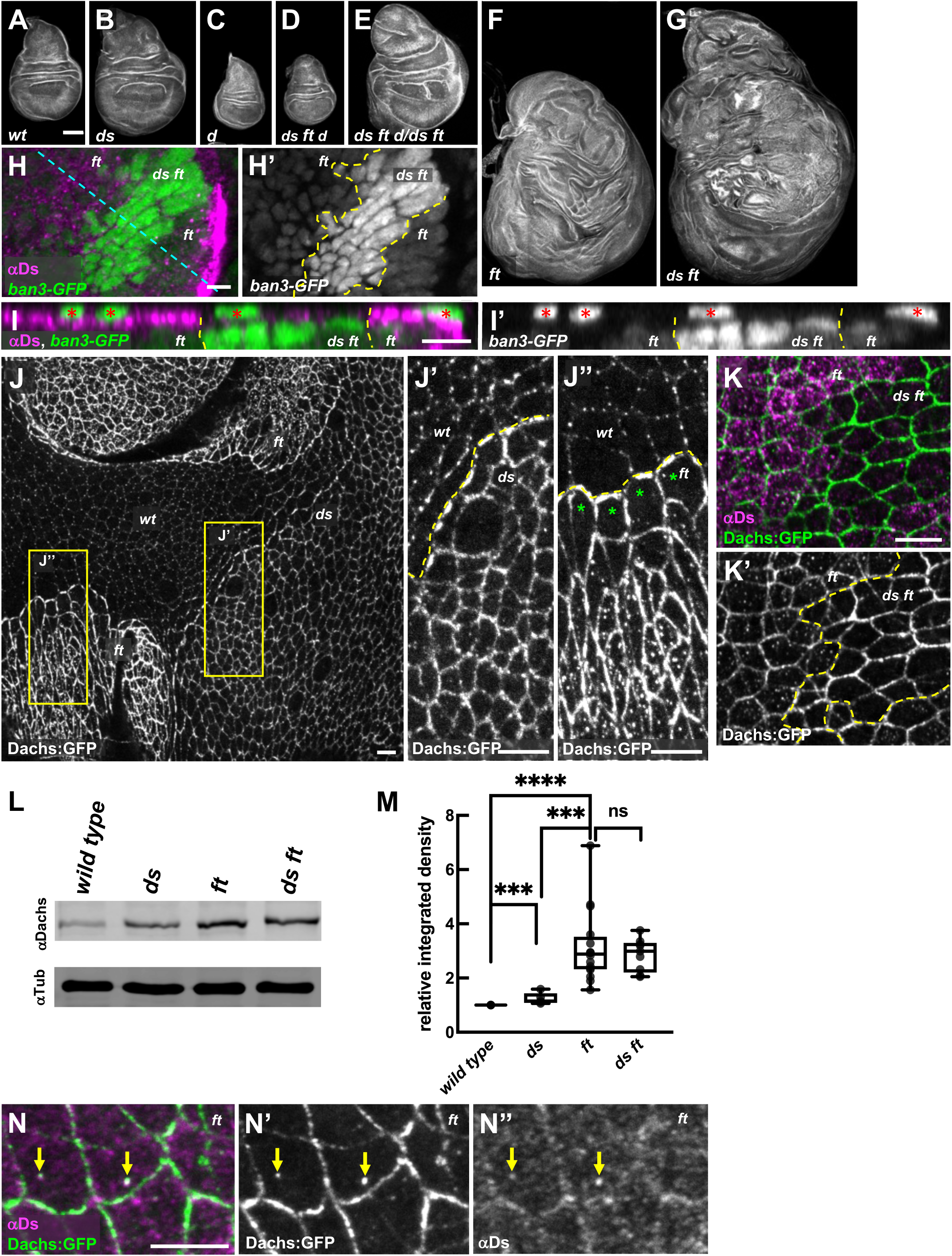
Both Ft and Ds suppress tissue growth by regulating localization and abundance of Dachs at the junctional cortex. (A-G) Representative wing discs stained with phalloidin showing the growth defects of the following genotypes: wild-type (A), *ds^UAO71^* (B), *dachs^GC13^* (C), *ds^UAO71^ ft^G-rv^ dachs^GC13^*/*ds^UAO71^ ft^fd^ dachs^GC13^* (D), *ds^UAO71^ft^G-rv^ dachs^GC13^*/*ds^UAO71^ ft^fd^* (E), *ft^G-rv^/ft^fd^* (F) and *ds^UAO71^ft^G-rv^*/*ds^UAO71^ft^fd^* (G). (H, I) Expression of *ban3-GFP*, a Hippo pathway reporter, compared between *ft^G-rv^/ft^fd^* (or *ds^UAO71^ ft^G-rv^*/*ft^fd^*) cells and *ds^UAO71^ ft^Gr-v^* cells in the wing disc. *ban3-GFP* is more highly expressed in *ds^UAO71^ ft^G-rv^* cells than in *ft* mutant cells. *ds ft* clones are marked by the absence of Ds staining. (I) XZ section indicated by the blue dashed line in H. Red asterisks indicate *ban3*-GFP expression in peripodial cells. (J) Comparison of Dachs:GFP levels and subcellular localization in wild type, *ds^UAO71^* and *ft^fd^* mutant cells. In *ds* mutant cells Dachs:GFP levels were slightly increased and primarily localized to the junctional cortex (J and J’). In *ft* mutant cells, Dachs:GFP levels increased more than in *ds* cells and localized both at the junctional cortex and to non-junctional puncta (J and J”). Clone boundaries and genotypes determined by anti−Ft staining (Fig. S1A). (K, K’) In comparison to *ft* cells, *ds ft* mutant cells displayed even more Dachs:GFP at the junctional cortex but lacked the non-junctional puncta. (L, M) Immunoblot analysis showing Dachs levels in imaginal tissue from wild-type, *ds^UAO71^*(N=5), *ft^G-rv^/ft^fd^* (N=16) and *ds^UAO71^ ft^G-rv^*/*ds^UAO71^ ft^fd^* (N=9). Dachs abundance increases slightly in *ds* but is much greater in *ft* and *ds ft* imaginal tissue. (M) Relative Dachs levels normalized to α-Tubulin. **** P<0.0001, *** 0.0001<P<0.001, ns not significant, (two-tailed unpaired t-test between each genotype). (N) Localization of Ds and Dachs:GFP in the absence of Ft. Dachs:GFP was highly localized to the junctional cortex and colocalized with Ds in non-junctional puncta. Scale bars: (A) 100 µm, (H-K, N) 5 µm. In this and all subsequent figures fluorescently-tagged proteins were viewed by native fluorescence. Antibody staining is indicated by an ’α’ symbol before the protein name.

To determine if Ds and Ft synergistically repress growth via the Hippo pathway, we compared the effect of *ds ft* double mutants to *ft* single mutant cells on the *ban3-GFP, a* reporter of Yki activity (Matakatsu and Blair, 2012). We observed that expression of *ban3-GFP* is upregulated to a greater extent in *ds ft* than in *ft* cells (Fig. 1H, I). Thus, Ds and Ft synergistically repress growth by repressing Yki activity.

### Ft and Ds restrict tissue growth by regulating the abundance of Dachs at the junctional cortex

The finding that Ft and Ds function synergistically to restrict growth seems at odds with the model that Ds promotes Dachs accumulation at the junctional cortex while Ft promotes its degradation. To explore this paradox, we examined Dachs localization and abundance in *ds ft* double mutants relative to *ds* or *ft* single mutants. Compared to wild type, we found that Dachs:GFP accumulated at slightly greater levels in *ds* mutant tissue (Fig. 1J, J’) and was even more abundant in *ft* mutant tissue (Fig. 1J). Additionally, we noticed that in *ft* mutant cells Dachs was distributed in puncta across the apical cortex except in mutant cells at the clone borders with wild-type cells (Fig. 1J, J’’). We infer this effect might occur because Ds protein in *ft* mutant cells at the edge of clones interacts with Ft and is tightly localized at the boundary with adjacent wild-type cells, while in the middle of *ft* clones Ds is mislocalized across the apical cortex (Strutt and Strutt, 2002; Ma et al., 2003; see below).

If Ds is necessary to recruit Dachs to the junctional cortex, then loss of both Ds and Ft should result in decreased Dachs cortical accumulation in comparison to loss of just Ft alone. Remarkably, we instead found that *ds ft* cells accumulated Dachs at the junctional cortex to even higher levels than those lacking Ft alone (Fig. 1K-K’, Fig. S1B-C). Furthermore, the non-junctional Dachs puncta we observed in *ft* mutant cells were absent in *ds ft* double mutant cells (Fig. 1K-K’). These results clearly indicate that neither Ds nor Ft is required for cortical localization of Dachs. Additionally, the increased level of cortical Dachs and absence of non-junctional Dachs puncta in *ds ft* double mutant cells suggests that Ds can promote removal of Dachs from the junctional cortex. Given that *ds ft* double mutant discs are significantly more overgrown than *ft* mutants alone (Fig. 1F, G, Matakatsu and Blair, 2006), the non-junctional Dachs we observed in *ft* mutant cells likely does not promote growth. To determine how loss of Ft and/or Ds affects Dachs abundance, we compared Dachs protein levels in *ds* null, *ft* null and *ds ft* null double mutant imaginal tissue using immunoblots (Fig. 1L, M). While *ds* mutant tissues displayed slightly higher levels of Dachs, presumably because loss of Ds causes decreased Ft activity (Sopko et al., 2009; Feng et al., 2009) Dachs abundance was significantly greater in *ft* mutant tissue than either *ds* or wild-type tissues.

However, in *ds ft* double mutant tissues, Dachs levels were similar to *ft* mutants (Fig. 1L, M), suggesting that increased cortical Dachs in *ds ft* double mutants relative to *ft* mutants (Fig. 1K) reflects changes in localization rather than abundance. These results also indicate that increased junctional accumulation of Dachs is critical to growth. Consistent with this idea, loss of one copy of *dachs* strongly suppressed *ds ft* overgrowth (Fig. 1E, G) and homozygous loss of *dachs* was epistatic to it (Fig. 1C, D, G).

### The Dachsous intracellular domain regulates growth in a context-dependent manner

Previous studies have shown that in the absence of Ft, Ds becomes mislocalized from the junctions and is much more diffusely distributed across the apical membrane (Strutt and Strutt, 2002; Ma et al., 2003; Fig. 1K, Fig. S1D). Because Ds and Dachs are known to physically interact (Bosveld et al., 2012), mislocalized Ds might recruit Dachs away from the junctional cortex, thereby reducing growth. Careful examination of Ds and Dachs in *ft* mutant cells revealed that both displayed non-junctional puncta that often co-localized (Fig. 1N), consistent with the possibility that Ds can remove Dachs from the junctional cortex.

To better understand how Ds affects Dachs subcellular localization, we next focused on the intracellular domain of Ds, which has been proposed to form a complex with Dachs and Dlish (Bosveld et al., 2012 and 2016; Zhang et al., 2016; Misra and Irvine, 2016). To dissect the function of Ds, we used CRISPR-Cas9 to replace the Ds ICD with GFP (Ds^ΔICD:GFP^; Fig. 2A). Ds^ΔICD:GFP^ expression levels and localization were indistinguishable from wild-type Ds (Fig. 2B, C and Fig. S1E, F). Ft also was normally localized in *ds^ΔICD:GFP^*cells, suggesting that Ds^ΔICD:GFP^ binds Ft thereby promoting its ability to reduce growth (Fig. 2D). In marked contrast to *ds* null wings and wing discs, which are slightly overgrown, *ds^ΔICD:GFP^* wings and wing discs were smaller than normal (Fig. 2E-K). This result is surprising given our evidence that Ds can repress growth by removing Dachs from the junctional cortex, but likely reflects the complex nature of Ds function in growth control: in the presence of Ft, Ds functions both to promote growth by binding and protecting Dachs from Ft-mediated degradation and to bind Ft thereby promoting its ability to reduce growth. If so, then loss of *ft* should be epistatic to *ds^ΔICD:GFP^*. Indeed, the *ds^ΔICD^ ft* double mutant combination produced severe overgrowth similar to that of *ds ft* double mutant combination (Fig. 2M, compare with *ds ft* double mutant in Fig. 1G). Taken together, our results indicate that the function of the Ds ICD in growth control is context-dependent: in the presence of Ft the Ds ICD promotes Dachs activity while in the absence of Ft the Ds ICD represses Dachs activity.

**Figure 2.**
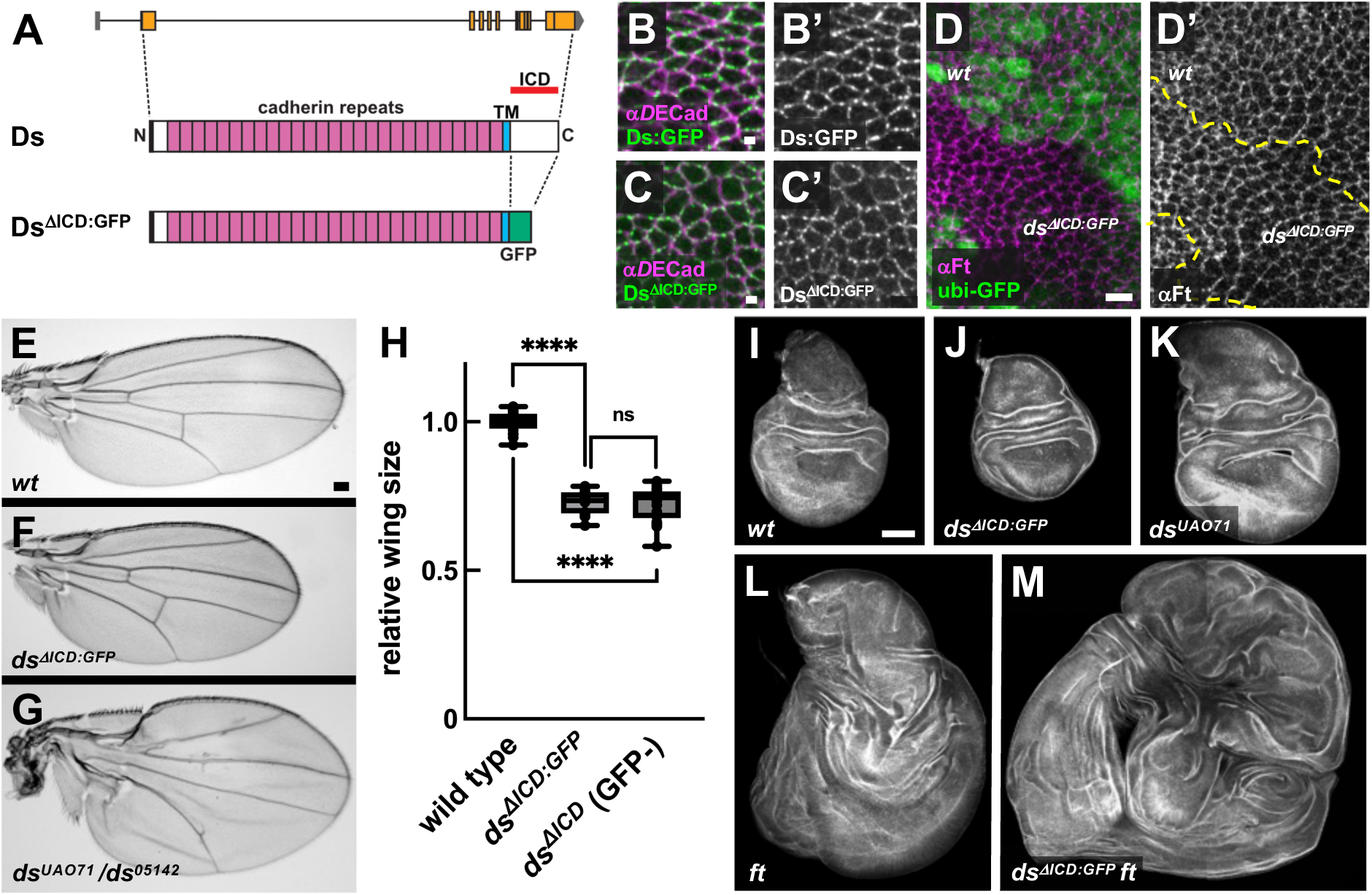
The Ds ICD has context-dependent effects on tissue growth. (A) Structure of Ds^ΔICD:GFP^. The intracellular domain was replaced with GFP (green) using CRISPR-Cas9. ICD, intracellular domain (red line) and TM, transmembrane domain (sky blue). (B and C) Localization of Ds^:GFP^ in a *ds^GFP^* homozygote (B) and Ds^ΔICD:GFP^ in a *ds^ΔICD:GFP^* homozygote (C). Ds:GFP and Ds^ΔICD:GFP^ localize at the junctional cortex. Adherens junctions are marked by *D*E-Cad staining. (D) Ft localization is normal in *ds^ΔICD:GFP^* mutant cells. Mutant clones are marked with the absence of nuclear GFP. (E-G) Representative adult wings in indicated genotypes. The *ds^ΔICD:GFP^* wing displays undergrowth (F). (H) Quantification of wing size in wild-type, *ds^ΔICD:GFP^* and *ds^ΔICD^* (GFP-) mutants. **** P<0.0001, ns not significant, (two-tailed unpaired t-test between each genotype). (I-M) Growth of wing discs with the indicated *ds* and *ft* alleles. Genotypes: wild type (I), *ds^ΔICD:GFP^* (J), *ds^UAO71^* (K), *ft^G-rv^/ft ^fd^* (L) and *ds^ΔICD:GFP^ ft^G-rv^*/*ds^ΔICD:GFP^ ft^fd^* (M). Although loss of the Ds ICD leads to undergrowth (J), in combination with loss of *ft* mutant it causes severe overgrowth (M). Scale bars, (B-C) 1 µm, (D) 5 µm, (E) 100 µm and (I) 10 µm.

### The intracellular domain of Dachsous organizes Dachs into cortical puncta

How does the Ds ICD regulate growth? Dachs and Dlish are known to co-localize with Ds in puncta at the junctional cortex in wing discs and to associate with the Ds ICD *in vitro* (Bosveld et al., 2012 and 2016; Brittle et al., 2012; Zhang et al., 2016; Misra and Irvine, 2016). Consistent with these studies, we found a strong correlation between wild-type Ds and Dachs in puncta at the junctional cortex (Fig. 3A, B). However, in *ds^ΔICD^*mutant clones, Dachs appeared less punctate and was irregularly localized at the junctional cortex (Fig. 3C). Moreover, unlike wild-type Ds, Ds^ΔICD:GFP^ did not strongly colocalize with Dachs (Fig. 3D-E).

**Figure 3.**
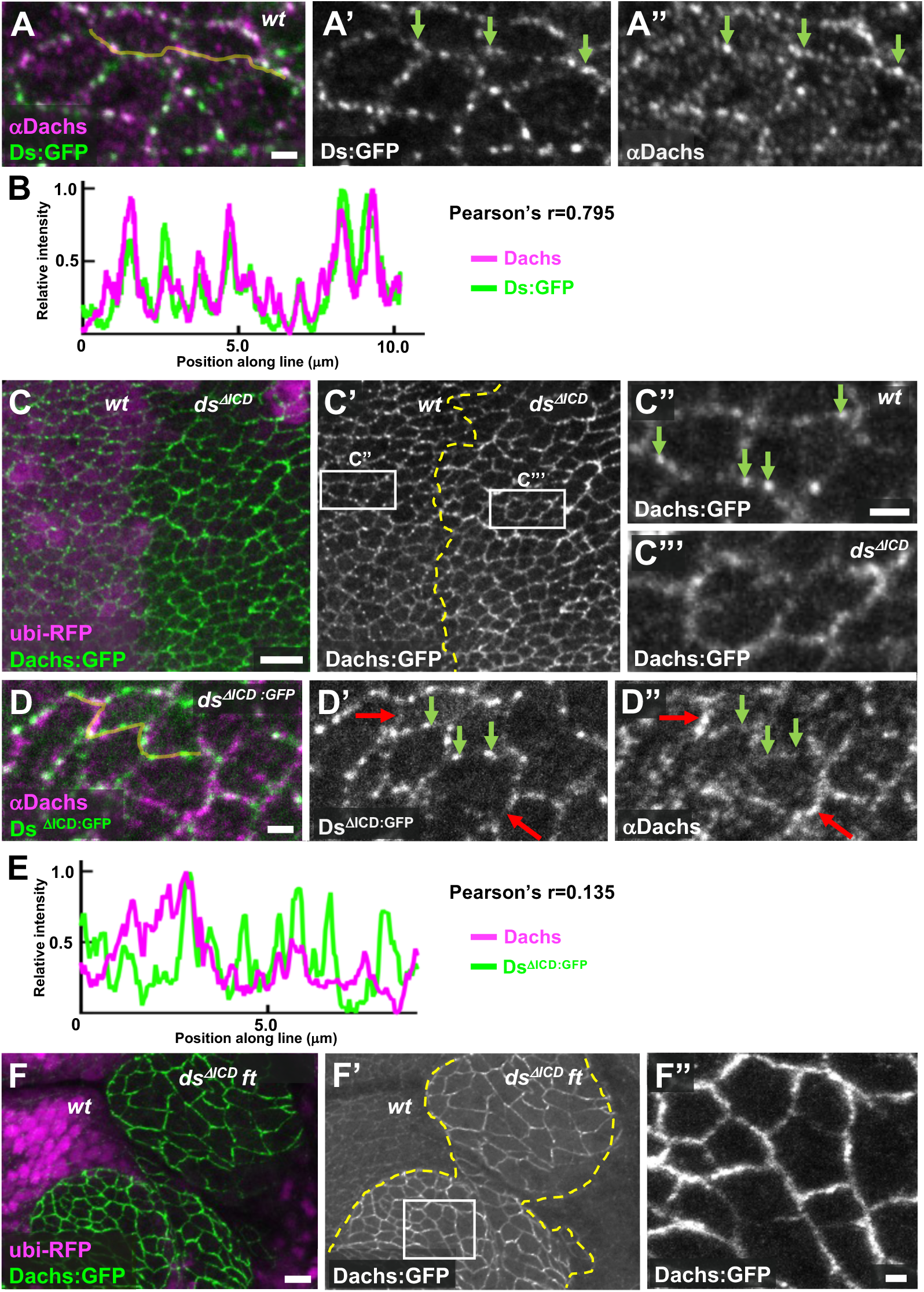
The Ds ICD influences Dachs localization. (A) Ds:GFP and Dachs colocalize at junctional puncta (green arrows) in wing discs. (B) Fluorescence intensity of Dachs and Ds:GFP along the yellow line in A showing strong colocalization. (C) In *ds^ΔICD^* clones, Dachs:GFP loses its punctate appearance and becomes more uniformly distributed along the junctional cortex (the absence of nuclear RFP serves as clonal marker). (D) Localization of Dachs and Ds^ΔICD:GFP^ in a *ds^ΔICD:GFP^* homozygote. Dachs fails to colocalize with Ds^ΔICD:GFP^ (green arrows indicate Ds^ΔICD:GFP^ puncta). (E) Fluorescence intensity of Dachs and Ds^ΔICD:GFP^ along the yellow line in D showing weak colocalization in comparison to B (Pearson’s correlation coefficient r=0.135 vs r=0.795). (F-F“) Dachs:GFP accumulates strongly and almost exclusively at the junctional cortex in *ds^ΔICD^ ft* mutant clones. Note the absence of non-junctional puncta of Dachs:GFP here in comparison to *ft* mutant cells in Figure 1J”. Scale bars, (A, C”, D, F”) 1 µm (C, F) 5 µm.

In the absence of *ft*, Dachs colocalizes with Ds in non-junctional puncta (Fig. 1J’’, N). To ask if the Ds ICD is necessary for this effect, we examined Dachs in *ds^ΔICD^ ft* double mutant tissue. We found that non-junctional Dachs puncta were dramatically reduced and instead Dachs accumulated strongly at the junctional cortex, similar to *ds ft* double mutants (Fig. 3F). Given that *ds^ΔICD^ ft* and *ds ft* double mutants display similar overgrowth phenotypes (Figs. 1G and 2M), these results suggest that the Ds ICD associates with Dachs, thereby influencing Dachs’ localization and growth-promoting function at junctional cortex.

### Dachsous regulates Approximated localization at the junctional cortex

App encodes a multi-pass transmembrane palmitoyltransferase that is thought to regulate both Ft and Dlish through post-translational palmitoylation (Matakatsu and Blair, 2008; Matakatsu et al., 2017; Zhang et al., 2016). App is also localized to the junctions (Matakatsu and Blair, 2008), but what determines its localization is unclear. To visualize App, we tagged it at the C-terminus with YFP (App:YFP) or in the extracellular domain with V5 (App:V5) using CRISPR-Cas9 (Fig. S1G-I). Both tagged alleles display normal wing and leg size and shape, indicating the tags do not interfere with App function. App:YFP localized to junctional puncta in a similar pattern to Dachs and Ds and was predominantly localized at proximodistal junctions in wing discs (Fig. 4A-D), reminiscent of Dachs, Dlish and Ds planar polarized distribution (Brittle et al., 2012; Ambegaokar et al., 2012; Misra and Irvine, 2016). In fact, App colocalized with Dachs, Ds and Dlish (Fig. 4A-D and Fig. S1J, see below).

**Figure 4.**
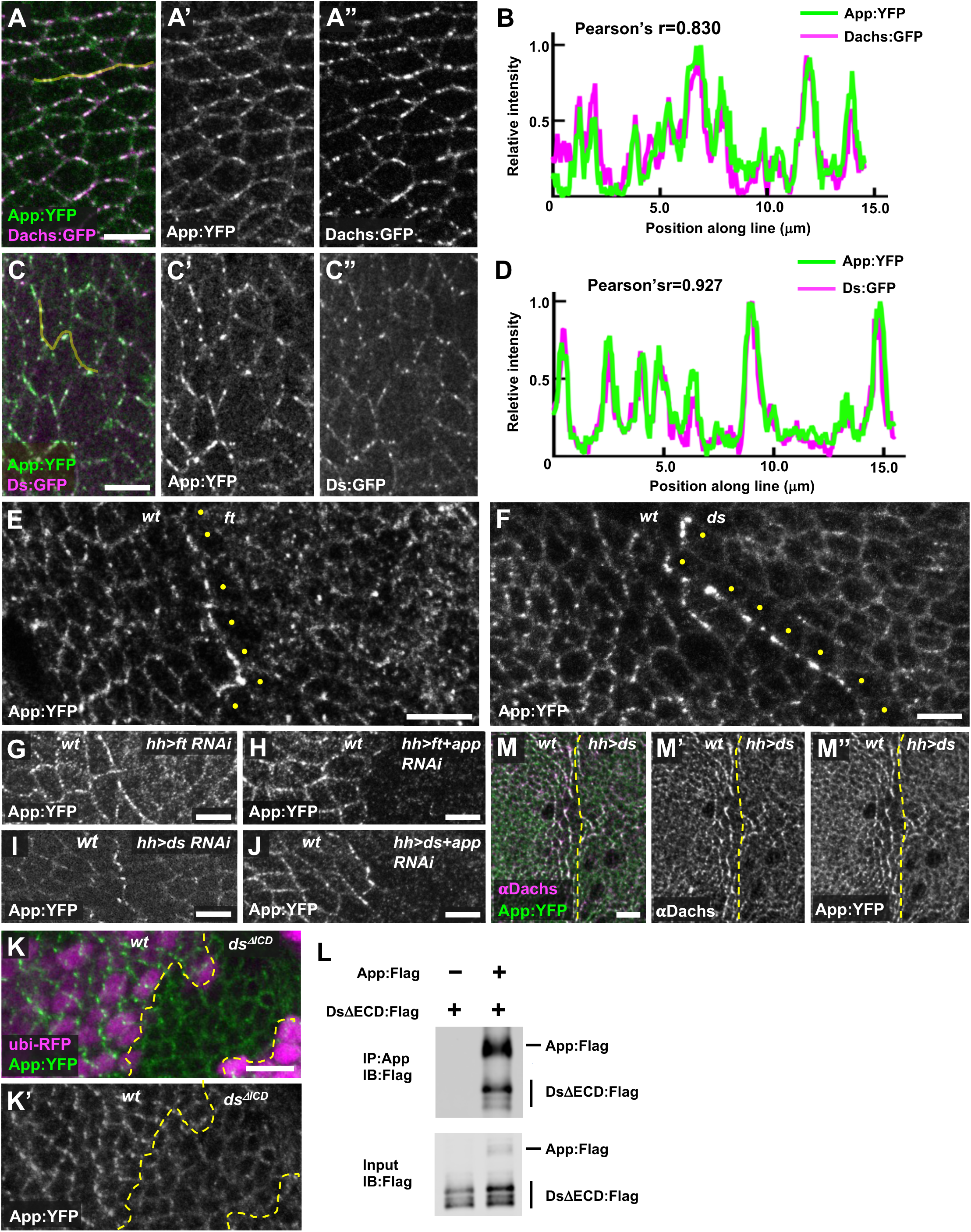
Ds and Ft regulate App at cell junctions. (A-D) Ds, Dachs and App colocalize at junctional puncta. Representative image (A) and line scan quantification of fluorescence intensity (B) showing colocalization of App:YFP and Dachs:GFP. Representative image (C) and line scan quantification of fluorescence intensity (D) showing colocalization of Ds:GFP and App:YFP. (E) Loss of *ft* causes mislocalization of App:YFP from junctional puncta to a more uniform distribution across the apical surface (clonal marker not shown). Note that App:YFP accumulates at the boundary between wild-type and *ft* mutant cells and appears reduced in the cytoplasm of *ft* cells at the edge of the clone (yellow dots). (F) The localization of App:YFP is altered in *ds* mutant cells. App:YFP is more diffuse in the absence of Ds and accumulates at the clone boundary (clonal marker not shown). Note that cytoplasmic App:YFP is reduced in *ds* cells adjacent to the boundary (yellow dots). (G, H) App:YFP accumulates at the boundary between wild-type and *ft* RNAi cells (G) but fails to accumulate at the boundary in *ft*, *app* double RNAi cells (H), indicating that App in *ft* depleted cells is recruited to contacts with wild-type cells. (I, J) App:YFP accumulates at the boundary between wild-type and *ds* RNAi cells (I) and at the boundary between wild-type and *ds, app* double RNAi cells (J), indicating that App in *ds* depleted cells is not recruited to contacts with wild-type cells. (K) Junctional puncta of App:YFP are much less apparent in *ds^ΔICD^* clones. (L) The Ds ICD co-immunoprecipitates App in S2 cells. (M) Ectopic expression of Ds in the posterior compartment causes Dachs and App:YFP to colocalize in a planar polarized fashion for 2-3 cell diameters outside the Ds overexpression domain. Scale bars, 5 µm.

We next explored App’s relationship with Ft and Ds. In *ft* mutant cells, App displayed decreased junctional accumulation but retained its punctate appearance, similar to Ds (Fig. 4E and Fig. S1D). In *ds* mutant cells, App displayed junctional localization and lacked distinct puncta (Fig. 4F). Notably, App distribution differed significantly from both Dachs and Dlish, which displayed increased accumulation at the junctional cortex in either *ft* or *ds* mutant cells (Fig. 1J, Bosveld et al., 2012; Brittle et al., 2012; Zhang et al., 2016; Misra and Irvine, 2016). These observations suggest that Dachs and Dlish are regulated differently from App and that App is more directly dependent upon Ds for junctional localization.

Previous studies have shown that due to their heterophilic interactions, Ft and Ds accumulate at the boundary of either *ds* or *ft* mutant clones (Strutt and Strutt, 2002; Ma et al., 2003). We found that App also accumulated at these boundaries (Fig. 4E, F), raising the possibility that Ft or Ds recruits App. To determine this, we used *ft* RNAi in the posterior compartment of the wing (driven by *hh-Gal4*) to create polarized accumulation of Ds on the posterior side of the boundary and Ft on the anterior (wild-type) side. Conversely, *ds* RNAi in the posterior caused Ft to accumulate on the posterior side of the boundary and Ds to accumulate on the anterior (wild-type) side. As we observed in somatic mosaic clones (Fig. 4E, F), App accumulated at the boundary between wild-type cells and either *ft* (Fig. 4G) or *ds* knockdown tissue (Fig. 4I). To determine on which side of the boundary App accumulated, we used double RNAi to deplete both App and either Ft or Ds. Co-depletion of Ft and App in the posterior strongly reduced accumulation of App at the boundary (Fig. 4H), indicating that App accumulates with Ds in the posterior (Ft-depleted) cells at the boundary. Conversely, co-depletion of Ds and App did not affect App accumulation at the boundary (Fig. 4J), indicating that when Ds is depleted App accumulates in adjacent wild-type cells at the boundary. In short, these results show that Ds recruits App to these boundaries while Ft does not.

Three further observations suggest that the Ds ICD interacts with App. First, App puncta were less apparent at the junctional cortex in *ds^ΔICD^* mutant clones (Fig. 4K). Second, App co-immunoprecipitated with the Ds ICD in S2 cells (Fig. 4L). Third, when we ectopically expressed Ds in the posterior compartment we found that as with Dachs, this resulted in repolarization and stabilization of App in cells just anterior to the compartment boundary (Fig. 4M, Bosveld et al., 2012; Brittle et al., 2012; Ambegaonkar et al., 2012). Taken together, our results are consistent with the idea that the Ds ICD coordinately recruits and stabilizes Dachs and App (and possibly Dlish) in puncta at the junctional cortex.

### Mapping functional domains in the Ds ICD

We next mapped functional domains in the Ds ICD using a combination of cell culture and *in vivo* experiments. To identify binding domains in the Ds ICD, we used a previously described approach to polarize protein domains at the cortex of cultured S2 cells (Johnston et al., 2009; Bosveld et al., 2016; Tokamov et al., 2023). Specifically, we fused the full-length or truncated Ds ICD to the extracellular domain of the homophilic adhesion protein Echinoid (Ed), which causes cells to aggregate and accumulate the fusion protein at points of contact. Using this approach, we asked which domains in the Ds ICD are necessary to recruit core complex proteins (Fig. 5A). In control cells, the Ed extracellular domain fused to Flag (Ed:Flag) accumulated at cell-cell contacts but co-transfected core complex proteins instead localized to the cytoplasm or to cytoplasmic foci (Fig. S2A). In contrast, Ed fused to Ds ICD (Ed:Ds:Flag) recruited core complex proteins to cell contacts (Fig. 5B, C).

**Figure 5.**
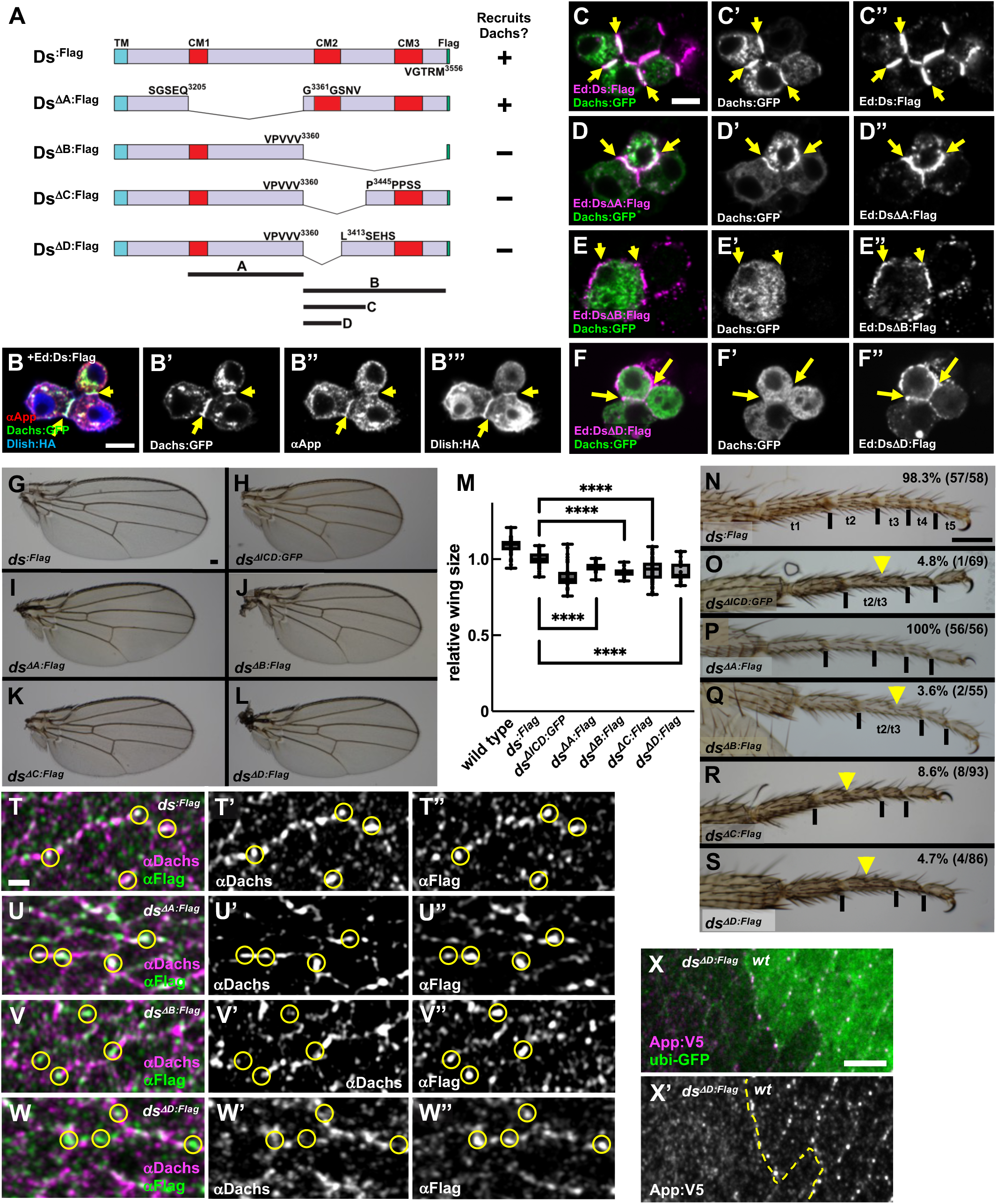
Mapping Ds functional domains. (A) Structure of the Ds ICD and the deletions generated for the S2 cell recruitment assay and CRISPR mutations *in vivo*. Full-length Ds ICD or Ds ICD deletions (ΔA,ΔB, ΔC and ΔD) were fused to the Ed extracellular domain for S2 cell experiments. The same domains are deleted with CRISPR-Cas9 *in vivo*. Abbreviations: Conserved motif 1-3 (CM1-3, red boxes): transmembrane domain (TM, sky blue box). (B) Ed:Ds:Flag recruits Dachs:GFP, App and Dlish:HA to cell contacts (yellow arrows). Ed homophilic interactions promote accumulation of the Ed fusion proteins at cell-cell contacts. (C-F) Recruitment of Dachs:GFP by Ds ICD and its deletions. Ed:Ds:Flag and Ed:DsΔA:Flag recruit Dachs:GFP to cell contacts (yellow arrows), but Ed:DsΔB:Flag and Ed:DsΔD:Flag fail to recruit Dachs:GFP. (G-L) Adult wings of the indicated genotypes. (M) Quantification of wing size in wild type and *ds* ICD deletion mutants. **** P<0.0001, (two-tailed unpaired t-test between each genotype). (N-S) Tarsal segments in legs from flies of the indicated genotypes. Numbers in the top right corner indicate the percentage of animals with normal tarsal segments. (T-W) Dachs localization in wing discs of the indicated genotypes. Dachs colocalizes with Ds in *ds^:Flag^* (T) and *ds^ΔA:Flag^* (U), but they colocalize less in *ds^ΔB:Flag^*(V) and *ds^ΔD:Flag^* (W). Yellow circles indicate Ds puncta. (X) App puncta, stained extracellularly with a pulse-chase protocol (see Methods) are lost in *ds^ΔD:Flag^* clones. Scale bars, (B, C, X) 5 µm, (T) 1µm, (G, N) 100 µm. In this and subsequent figures where epitope tags (Flag, HA, V5) are indicated antibodies were used that recognize those tags.

To determine the essential domains for core complex recruitment by Ds ICD, we first focused on two non-overlapping domains we denote A and B (Fig. 5A), because each contains <u>c</u>onserved <u>m</u>otifs (CM1-3) found in mammalian Ds (Hulpiau and Roy, 2009). Core complex proteins were recruited to cell contacts by domain B (Ed:Ds ΔA:Flag; Fig. 5D), but not by domain A (Ed:Ds ΔB:Flag; Fig. 5E). We then defined a smaller region within B (domain D) that includes CM2 and is necessary for recruitment (Fig. 5A, F, Fig. S2B-E).

To examine the functional significance of these domains *in vivo*, we generated fly lines lacking these portions using CRISPR-Cas9 (Fig. 5G-S). Interestingly, *ds^ΔA:Flag^*, *ds^ΔB:Flag^, ds^ΔC:Flag^* and *ds^ΔD:Flag^* all led to undergrowth in the wing that was milder than that of *ds^ΔICD:GFP^* (Fig. 5G-M). Additionally, we noted that *ds^ΔB:Flag^, ds^ΔC:Flag^* and *ds^ΔD:Flag^* displayed defects in leg tarsal segmentation commonly observed in Ft signaling mutants while *ds^ΔA:Flag^*did not (Fig. 5N-S). Thus, while both domain A and B are involved in growth control, domain B alone is sufficient for the Ds-Ft signaling functions that pattern the leg.

To determine if results from our S2 cell experiments reflect *in vivo* functions, we asked if Ds deletions recruit the core complex to junctional puncta (Fig. 5T-X). The expression pattern, protein level and subcellular distribution of all Ds mutant proteins were indistinguishable from endogenous Ds or Ds:Flag (Fig. S3A-D). While Dachs colocalized with Ds:Flag and DsΔA:Flag at junctional puncta (Fig. 5T-U, Fig. S3E-F), it did not colocalize as well with DsΔB:Flag and DsΔD:Flag (Fig. 5V-W, Fig. S3G-H). Moreover, App was dramatically reduced in *ds^ΔB:Flag^* and *ds^ΔD:Flag^* (Fig. 5X and data not shown). Taken together, our results suggest that domain D is responsible for core complex recruitment to the junctional cortex while domain A might control growth in parallel through other effectors (see discussion). Consistent with this idea, animals carrying the heteroallelic combinations *ds^ΔA:Flag^*/*ds^ΔB:Flag^* or *ds^ΔA:Flag^/ds^ΔD:Flag^* showed normal tissue growth in the wing and leg patterning (Fig. S3I-N).

### The Fat intra- and extracellular domains have opposing functions

In the absence of Ft, we propose that Ds, which is no longer anchored junctionally, represses growth by removing Dachs from the junctional cortex. If so, then restoring Ds junctional localization, without restoring Ft’s ability to degrade Dachs, should strongly enhance tissue growth. To do this, we deleted the Ft ICD using CRISPR-Cas9 (*ft^ΔICD^*; Fig. S4A). Interestingly, we observed much greater overgrowth in *ft^ΔICD^* wing discs compared to *ft* null mutant tissues (Fig. 6A-C). Consistent with the idea that Ft^ΔICD^ promotes growth by holding Ds (and Dachs) at the junctional cortex, we found that in *ft^ΔICD^* mutant cells Ds was localized in an almost normal pattern (Fig. 6D), in striking contrast to Ds in *ft* null mutant tissue (Fig. 1K, Fig. S1D). Additionally, although Dachs was mislocalized in non-junctional puncta in *ft* null mutant cells (Fig. 1J, K), Dachs was predominantly localized to the junctional cortex in *ft^ΔICD^* mutant cells (Fig. 6D). App appeared normally localized at the junctional cortex in *ft^ΔICD^* mutant cells (Fig. 6E), in striking contrast to *ds* or *ds^ΔICD^* mutant cells (Fig. 4F, K).

**Figure 6.**
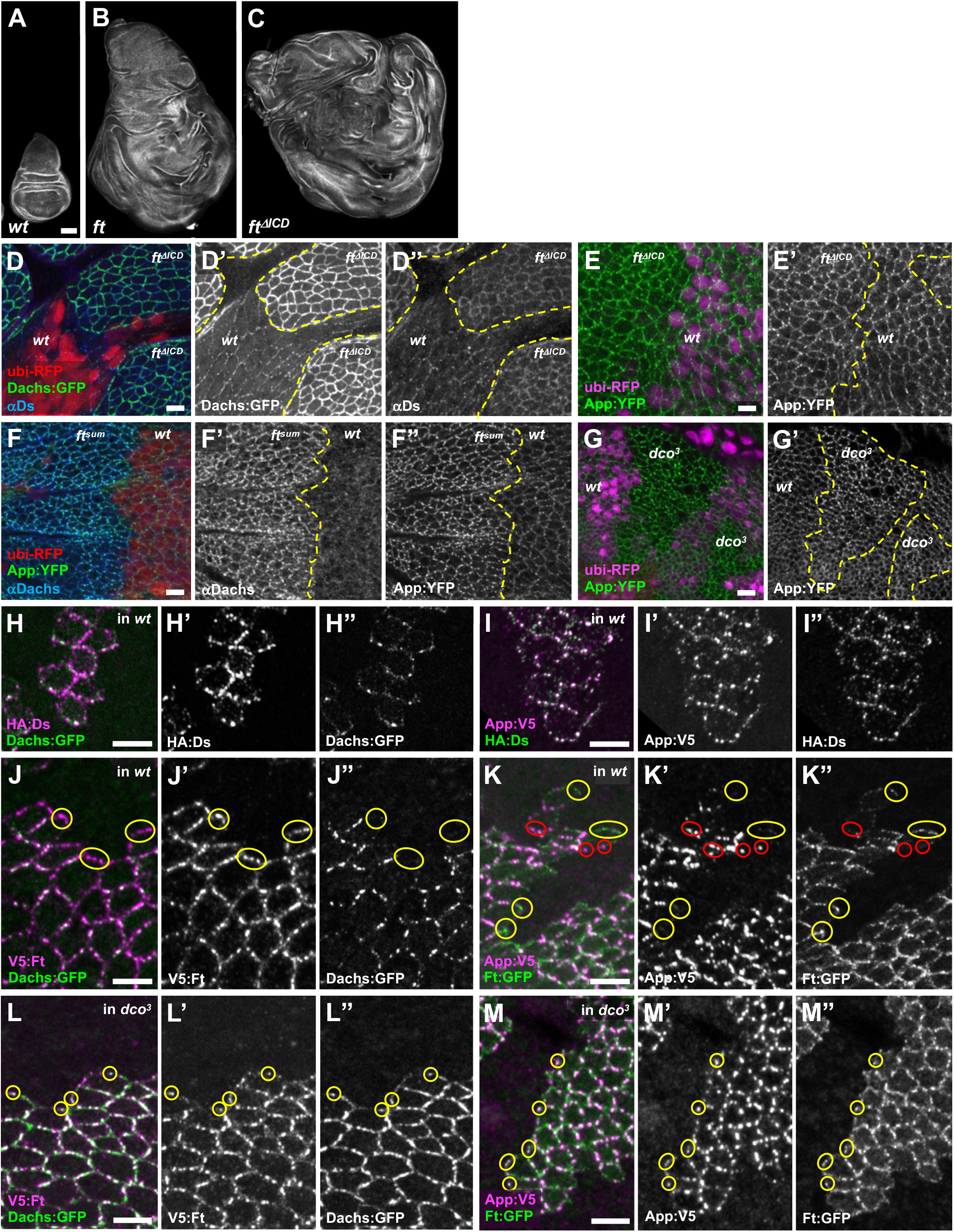
Effects of the Ft ICD on Dachs and App. (A-C) Representative wing discs of the following genotypes stained with phalloidin: wild-type (A), *ft^G-rv^/ft^fd^* (B) and *ft^ΔICD^* (C). Note that *ft^ΔICD^* has a more severe overgrowth phenotype than *ft* null mutants. (D) Localization of Dachs and Ds in *ft^ΔICD^*mutant cells. Dachs:GFP expression is elevated and uniformly distributed around at the junctional cortex, while Ds is only slightly elevated and localizes more normally at the junctions. Note that Dachs does not display the non-junctional puncta that are seen in *ft* null mutant cells (Fig. 1 J”). (E) Localization and abundance of App:YFP is not altered in *ft^ΔICD^* mutant cells. (F) Dachs and App:YFP levels are elevated but localization to junctional puncta does not change in *ft^sum^* mutant cells. (G) App:YFP abundance also is elevated at junctional puncta in *dco^3^*mutant cells. (H-M) Surface labeling of cell clones expressing HA:Ds (H, I), App:V5 (I, K, M) or V5:Ft (J, L) together with the indicated tagged proteins. All clones shown border non-expressing sister clones. (H) Dachs:GFP puncta colocalize with HA:Ds at clone boundaries. (I) Similarly, HA:Ds in puncta colocalizes with App:V5. (J) In contrast, V5:Ft puncta do not colocalize with Dachs:GFP at clone boundaries (yellow ellipses). (K) Similarly, Ft:GFP puncta do not co-stain for App:V5 at clone boundaries (yellow ellipses). Additionally, Ft:GFP fails to co-localize with App:V5 puncta at clone boundaries on the opposite side of cells (red ellipses). (L) In contrast to wild type, in a *dco^3^* mutant background V5:Ft and Dachs:GFP colocalize in puncta at clone boundaries (yellow circles). (M) Similarly, in a *dco^3^* mutant background App:V5 and Ft:GFP extensively colocalize in puncta at clone boundaries (yellow ellipses). Scale bars, 100 µm (A), 5 µm (D-M)

To further identify functions of the Ft ICD, we examined the effects of mutations that severely diminish activity of the Ft ICD without affecting its extracellular interactions with Ds. *ft^sum^* and *ft^61^* previously were identified as point mutations in the Ft ICD that cause overgrowth (Bossuyt et al., 2014; Bosch et al., 2014). Examining *ft^sum^* and *ft^61^*mutant clones, we found increased junctional accumulation of both Dachs and App in a punctate pattern (Fig. 6F and Fig. S4B, C). These observations are in contrast to *ft^ΔICD^* mutant clones, where App levels were unaffected (Fig. 6E) and Dachs was evenly distributed along the junctional cortex (compare Fig. 6D-F). Clones of *dco^3^* (discs-overgrown) cells, which also decrease activity of the Ft ICD by blocking its phosphorylation (Sopko et al., 2009; Feng and Irvine, 2009), had a similar effect on App (Fig. 6G). Interestingly, Dco fails to promote phosphorylation of Ft^sum^ in S2 cells (Fig. S4D), suggesting that phosphorylation of Ft decreases core complex accumulation at junctional puncta.

While the Ft ICD clearly has a role for reducing core protein levels, there is evidence that it recruits core proteins to the junction before degrading them. In fact, the Ft ICD forms a complex with core complex proteins including Dlish and App in S2 cells (Matakatsu et al., 2017; Zhang et al., 2016; Misra and Irvine, 2016). To further examine this possibility, we made use of the planar polarized distribution of Ft and Ds in the imaginal epithelium, where Ft accumulates primarily at the proximal junctions of cells and Ds accumulates primarily at the distal edge. By examining mosaic clones of cells expressing tagged forms of Ds, Ft, Dachs or App, we could discern at which junction, distal or proximal, these proteins accumulate in either wild-type or *dco^3^* backgrounds. As previously suggested, we found that Ds and Dachs accumulated at the distal junctions of wild-type imaginal epithelial cells (Fig. 6H), while Ft accumulated on the proximal side (Fig. 6J-K, Mao et al., 2006; Rogulja et al., 2008; Brittle et al., 2012; Bosveld et al., 2012). Additionally, although App colocalized with Ds at distal junctions (Fig. 6 I), Ft did not colocalize with App on proximal junctions suggesting that App localized distally with the rest of the core complex (Fig. 6K). In contrast, when similar clones were generated in the *dco^3^* mutant background, Dachs and App colocalized with Ft at the proximal junctions (Fig. 6L, M). Taken together these observations suggest that like Ds, Ft recruits App and Dachs to the junctional cortex but prevents their retention by promoting their degradation.

### Mapping functional domains in the Ft ICD

To further examine the functions of the Ft ICD, we employed the Echinoid S2 cell assay to examine the ability of Ed:Ft ICD fusions to recruit core complex components to the cell cortex (Fig. 7A). Just as we observed with Ds ICD fusions, Ed:Ft:Flag strongly recruited Dachs, App and Dlish (Fig. 7A). Recruitment of Dachs did not appear to depend on other core complex proteins, as RNAi-mediated depletion of Dlish and App did not affect Dachs recruitment (Fig.7B). Similarly, we found that Dachs co-immunoprecipitated with the Ft ICD and that depletion of App and Dlish did not affect their co-immunoprecipitation (Fig. 7C). These results suggest that while the Ft ICD can recruit Dachs, Dlish and App to the cortex of S2 cells, it likely cannot promote their degradation, presumably because other proteins essential for this process are not expressed or active in S2 cells.

**Figure 7.**
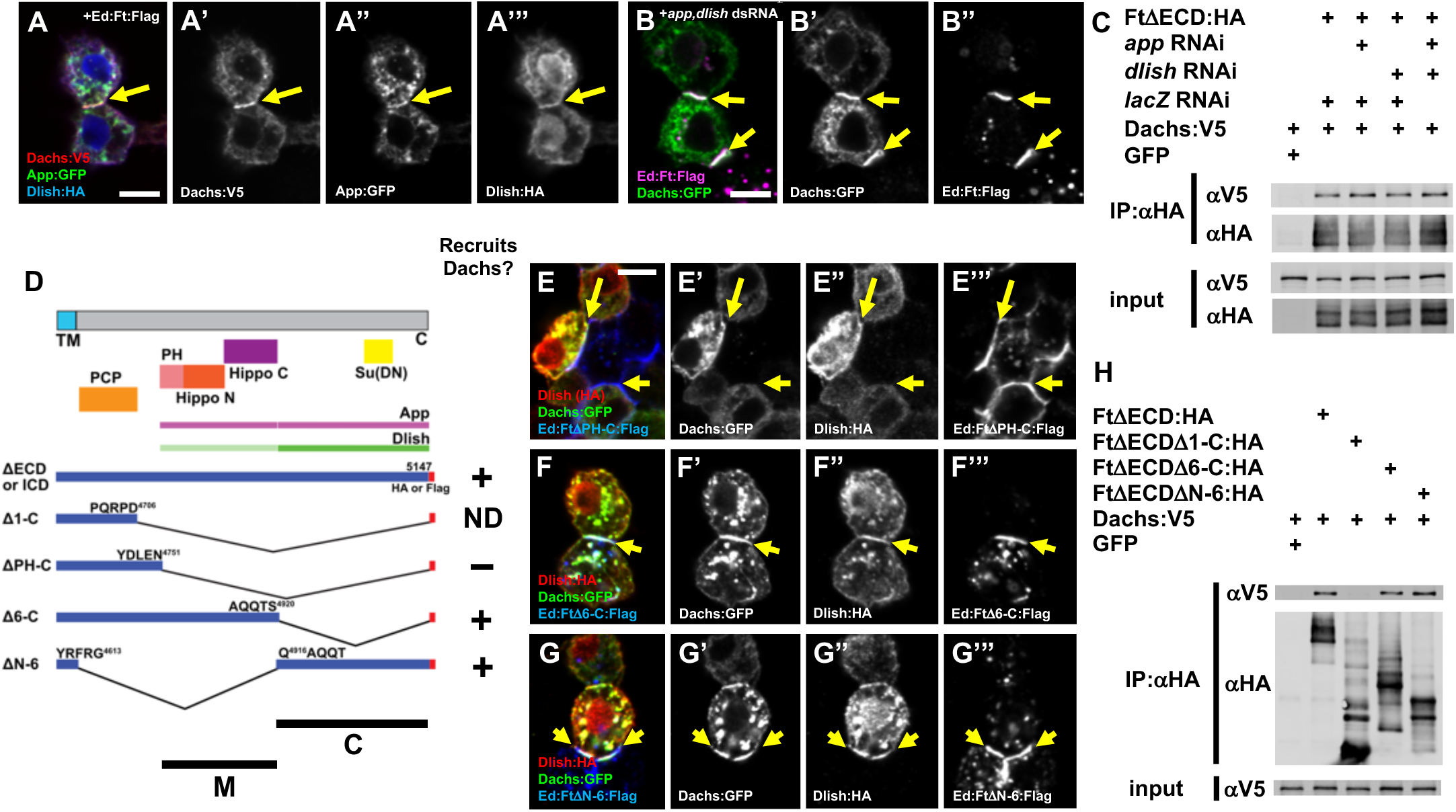
Interactions between the Ft ICD and core complex proteins in S2 cells. (A) Recruitment of Dachs:V5, App:GFP and Dlish:HA by Ed:Ft:Flag in S2 cells. All three proteins accumulate with the Ft ICD at cell contacts (yellow arrows). (B) Depletion of *app* and *dlish* by RNAi does not affect the ability of the Ft ICD to recruit Dachs:GFP (yellow arrows). (C) Co-immunoprecipitation experiments similarly show that the Ft ICD forms a complex with Dachs in S2 cells even when App and Dlish are depleted. (D) Structure cartoons of Ft ICD deletions designed to identify regions responsible for core complex recruitment in S2 cells. Functional domains and App and Dlish association sites from previous studies are also shown. ND: not determined. (E-G) S2 cell assays for recruitment of core complex proteins by different Ft ICD deletions. (E) Ed:FtΔPH-C:Flag fails to recruit core complex proteins to cell contacts. Ed:FtΔ6-C:Flag (F) as well as Ed:FtΔN-6:Flag (G) recruit Dachs:GFP and Dlish:HA (yellow arrows). (H) Co-immunoprecipitation experiments confirm the results in E-G. Scale bars, 5 µm.

Previous studies have revealed the domains in the Ft ICD responsible for PCP and growth control *in vivo* (Fig. 7D, Matakatsu and Blair, 2012). To better understand how different Ft ICD domains function, we next examined their role in core complex recruitment (Fig. 7D-H). Ed:FtΔPH-C:Flag, which lacks most of the ICD but retains a domain known to function in planar cell polarity (Matakatsu and Blair, 2012), failed to recruit any of the core complex proteins (Fig. 7E). To narrow down regions necessary for recruitment, we generated two smaller deletions, one containing the Hippo N and C domains previously shown to be sufficient for growth control (Ed:FtΔ6-C:Flag) and one that deletes those domains but retains regions that modulate Hippo pathway function (Ed:FtΔN-6:Flag; Matakatsu and Blair, 2012; see Fig. 7D). Interestingly, we found that either construct could recruit core complex proteins, consistent with previous work showing that both the M and C domains co-immunoprecipitate with App and Dlish in S2 cells (Fig. 7F-H, Zhang et al. 2016; Misra and Irvine, 2016).

### The relationship between Dachs, Dlish and App

The different effects of loss of Ft and Ds on Dachs, Dlish and App led us to further explore the relationship between these proteins. In the simplest case, these three proteins act together in a complex to promote growth. Consistent with this idea, Dachs interacts with Dlish *in vitro*, ectopic Dachs expression strongly stabilizes Dlish at the junctional cortex, and vice versa (Zhang et al., 2016; Misra and Irvine, 2016). Additionally, loss of App disrupts cortical localization of Dachs and Dlish (Matakatsu and Blair, 2008; Matakatsu at el., 2017; Zhang et al., 2016; Misra and Irvine, 2016). However, the relationship between Dachs or Dlish and App at the junctional cortex is less well studied. To address this, we expressed Dachs or Dlish and found that either promotes accumulation of App at the junctional cortex (Fig. 8A, C and Fig. S5A). In contrast, Dachs and Dlish levels changed only slightly in response to ectopic App expression, though they did have a more punctate appearance (Fig. 8B, C, Fig. S5B). Thus, while ectopic Dachs or Dlish can promote App accumulation, ectopic App has little effect on Dachs or Dlish.

**Figure 8.**
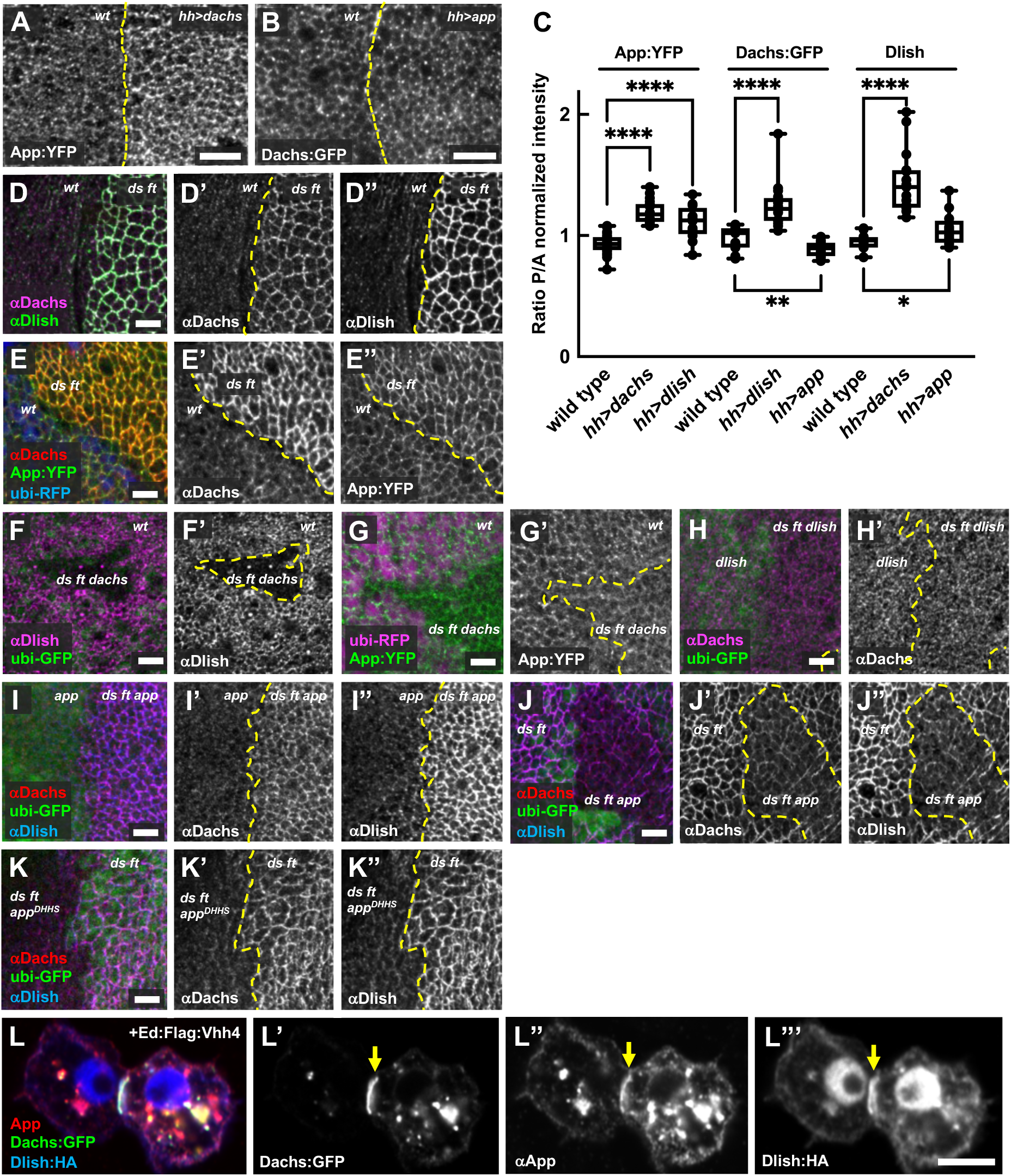
Dachs, App and Dlish form a core complex to promote growth. (A) Ectopic Dachs expression causes increased App abundance. (B) Conversely, ectopic App expression does not increase Dachs abundance but does result in a more punctate appearance at the junctional cortex. (C) Comparison of apical protein levels of Dachs, Dlish and App in response to ectopic expression of other core complex components. Y axis shows the mean intensity ratio of the posterior compartment (ectopic expression) to the anterior compartment (normal cells). **** P<0.0001, ** 0.001<P<0.01, * 0.01<P<0.05, (two-tailed unpaired t-test between each genotype). (D-K) Analysis of core complex components at the cell cortex in the absence of Ds and Ft. (D) Dachs and Dlish levels are highly elevated at the junctional cortex in *ds ft* mutant cells. (E) App:YFP abundance also is increased at junctions together with Dachs in *ds ft* clones. (F) In marked contrast to *ds ft* mutant cells (D’’), Dlish is strongly reduced in *ds ft dachs* mutant cells. (G) App:YFP is less punctate, but is junctionally localized in *ds ft dachs* mutant cells. (H) In the absence of *dlish*, Dachs abundance and localization are unaffected by loss of *ds* and *ft*. (I) In contrast, loss of *ds* and *ft* still causes increased Dachs and Dlish abundance at the junctional cortex in the absence of *app*, although the effect is smaller than when App is present (J). (K) Mutation in App catalytic domain has a similar effect on Dachs and Dlish accumulation as *app* null alleles (compare J and K). (L) Dachs:GFP localized to cell contacts also recruits App and Dlish in S2 cells. Scale bars, 5 µm

Although Ft and Ds affect accumulation of Dachs, App and Dlish at the junctional cortex, all three proteins accumulate at very high levels in the complete absence of *ds* and *ft* (Fig. 8D, E). Moreover, loss of *dachs* is epistatic to the *ds ft* overgrowth phenotype (Fig. 1D) and loss of either *dlish* or *app* partly suppresses this phenotype (Fig. S5C-I), indicating that these proteins can function together, independent of Ds and Ft, to promote growth. Therefore, we reasoned that it should be easier to understand how these proteins regulate their cortical accumulation and promote growth in *ds ft* mutants.

We started by examining Dlish and App in *ds ft dachs* triple mutant cells. In this background, Dlish was severely reduced from apical cortex (Fig. 8F), while App appeared more diffuse but still largely junctional (Fig. 8G). Loss of Dlish blocked the strong accumulation of Dachs normally seen in *ds ft* mutant cells (Fig. 8H). Taken together, these data suggest that Dachs and Dlish are strongly interdependent for their localization to the junctional cortex, while App can localize junctionally independent of Dachs and Dlish. However, we note that in *ds ft dachs* triple mutant cells App abundance is reduced relative to *ds ft* cells (compare Fig. 8E and G), suggesting that levels of Dachs and Dlish can affect App.

To explore the role of App in junctional recruitment of Dachs and Dlish we next examined these proteins in *ds ft app* triple mutant cells, either by removing *ds* and *ft* in an *app* mutant background (Fig. 8I) or by removing *app* in *ds ft* mutant background (Fig. 8J). Loss of *app* in *ds ft* background showed that App promotes accumulation of Dachs and Dlish at the junctional cortex in the absence of Ft and Ds (Fig. 8J), while loss of *ds* and *ft* in an *app* mutant background clearly showed that App is not absolutely required for Dachs and Dlish accumulation (Fig. 8 I). These observations are consistent with previous evidence that App palmitoylates Dlish *in vitro* and promotes junctional localization of Dlish as well as Dachs *in vivo*, since palmitoylation can promote association of cytoplasmic proteins with the plasma membrane (Zhang et al., 2016). To test this more directly, we examined the effect of *app^DHHS^*, a CRISPR engineered allele that blocks the palmitoyltransferase activity of App (Matakatsu et al., 2017), on Dachs and Dlish accumulation and found it to be indistinguishable from an *app* null allele (Fig. 8K). These results suggest that App’s primary role is to promote the accumulation of Dachs and Dlish at the junctional cortex through its enzymatic function rather than a structural one.

To examine interactions between core complex components, we modified the Echinoid S2 cell assay to work with any GFP-tagged protein by fusing an anti-GFP nanobody (Caussinus et al., 2012) to the cytoplasmic domain of Echinoid (pMT-Ed:Flag:vhh4) or to a myristoylation sequence (UAS-myr:Flag:vhh4). Ed:Flag:vhh4 expressed in S2 cells promoted cell aggregation and strongly accumulated at cell contacts, as shown previously for other Ed protein fusions (Johnston et al., 2009), and recruited co-transfected GFP (Fig. S5J). Ed:Flag:vhh4 strongly recruited Dachs:GFP to sites of cell contact, which in turn recruited co-transfected App and Dlish (Fig. 8L). This result was confirmed using myr:Flag:vhh4 (Fig. S5K). Additionally, Dlish and App could recruit other core complex components when they were tagged with GFP (Fig. S5L-O) in S2 cells. Thus, interactions among Dachs, Dlish and App help core complex formation in S2 cells.

## Discussion

In this study we have examined functional interactions between the intracellular domains of Ds and Ft and their effectors in growth control, a ‘core complex’ consisting of Dachs, Dlish and App. Previous work has suggested that while Ds functions to recruit core complex components to the cell cortex, thereby promoting growth, Ft promotes degradation of Dachs, thereby repressing growth (Mao et al., 2006; Bosveld et al., 2012; Brittle et al., 2012). The results presented here revise this model in several important ways. First, we show that the core complex requires neither Ft nor Ds to associate with the junctional cortex and drive growth. Rather, Ft and Ds function coordinately to precisely restrict the amount and distribution of core complex components at the junctional cortex. Second, we show that in the absence of Ft, Ds functions to repress growth, apparently by removing core complex components from the junctional cortex.

This finding is surprising and suggests that in some tissue contexts Ds could play a significant role in restricting tissue growth. Third we demonstrate that Ft also can recruit core complex components to the junctional cortex and that the stability and/or duration of these interactions is controlled by phosphorylation of the Ft ICD. Our structure/function analysis of Ft and Ds shows that they interact with core complex components via specific regions of their ICDs and that different regions of the Ds ICD function in parallel to promote growth. Taken together, our work reveals the underlying mechanisms by which Ds-Ft signaling represses a default ‘on’ state that is driven by the inherent ability of core complex components to localize to the junctional cortex and promote growth.

### The Dachs-Dlish-App core complex localizes at the junctional cortex to promote growth independent of Ft and Ds

Previous studies have found a strong correlation between the ability of Dachs to localize to the junctional cortex and its ability to promote tissue growth (Mao et al., 2006, Matakatsu and Blair, 2008, Zhang et al., 2016; Misra and Irvine, 2016). The results presented here strongly support this model. In particular, we find that although overall Dachs abundance is indistinguishable between *ft* mutant tissues and *ds ft* mutant tissues, Dachs abundance at the junctional cortex and overall tissue growth are much greater in the latter than in the former. Additionally, the finding that Dachs, Dlish and App all strongly accumulate at the junctional cortex in *ds ft* double mutant tissues demonstrates that core complex proteins accumulate at the junctional cortex independently of Ft and Ds.

To better understand how the core complex associates with the cell cortex, we examined interdependencies between these proteins in the absence of either Ft or Ds. Dachs and Dlish are highly interdependent - removing one results in almost complete loss of the other from the cell cortex - consistent with previous studies showing that Dachs and Dlish form a complex (Zhang et al., 2016; Misra and Irvine, 2016). App has also been shown to form a complex with Dachs and Dlish and we showed here that Dachs and Dlish can recruit App in S2 cells, but its relationship with these proteins at the cortex appears less direct. For example, while ectopic expression of Dachs or Dlish causes increased accumulation of App, ectopic expression of App has little effect on either Dach or Dlish. Additionally, loss of App reduces but does not prevent increased accumulation of Dachs and Dlish caused by absence of Ft and Ds. These observations are consistent with a model where App promotes the interaction of Dachs and Dlish with the cell cortex but does not contribute structural support for this interaction. Consistent with this idea, App can palmitoylate Dlish (Zhang et al., 2016) and we found that a mutation that specifically affects App enzymatic activity (*app^DHHS^*) affects Dachs and Dlish recruitment similar to an *app* null allele. Nonetheless, we consider App to be a member of the core complex because 1) co-IP and the Ed recruitment assays in S2 cells demonstrate that all three proteins can form a complex and 2) loss of Dachs severely diminishes App accumulation at the cell cortex (Matakatsu et al., 2017).

### Ds effects on growth are context dependent

Previous studies have highlighted the importance of Ds in stabilizing Dachs accumulation at the junctional cortex to promote growth (Bosveld et al., 2012; Brittle et al., 2012). While our findings are consistent with this model, they also show that in some contexts Ds can function to remove Dachs from the junctional cortex and as a result inhibit growth. This aspect of Ds function is clearly observed in the absence of Ft. As previously described (Hale et al., 2015), we observed that loss of Ft results in dramatic mislocalization of Ds, suggesting that Ds-Ft heterophilic interactions stabilize both proteins at the junctional cortex. We observed Ds and Dachs in non-junctional puncta in the absence of *ft.* These puncta were the consequence of Ds being unable to bind Ft on adjacent cells rather than a cell autonomous effect of loss of Ft because *ft* mutant cells at the borders of clones, where Ds could interact with Ft expressed on adjacent wild-type cells, did not display non-junctional Ds or Dachs. Additionally, these puncta were absent in in *ds^ΔICD^ ft* cells indicating that interaction with the Ds ICD is necessary to remove Dachs from the junctional cortex. Given that Ds is a transmembrane protein, it is possible that the non-junctional puncta are endocytic, but further studies will be required to address this.

What is most intriguing about these observations is that they provide a cell biological basis for the observation that while Ds promotes growth in the presence of Ft, it represses growth in the absence of Ft. While Ft is believed to play the major role in repressing Dachs activity and *ft* mutants display significant overgrowth, *ds ft* double mutants display much greater overgrowth. These observations raise the still unanswered question of whether, in some contexts, Ds might function to repress growth, independent of its role in activating Ft, during normal development. In relation to this it is interesting that the opposing gradients of Ds and Fj expression in the wing result in less stable Ds-Ft binding in the proximal wing (Hale et al, 2015). Further studies will be required to elucidate to what extent Ds represses growth independent of Ft in development.

### The Ft ICD can recruit the core complex to the cell cortex

Although Ft is well recognized to promote Dachs degradation, the underlying mechanistic basis of this function is poorly understood. Using the Echinoid S2 cell assay, we found that the Ft ICD functions similarly to the Ds ICD in having the ability to recruit core complex components to the cell cortex, consistent with previous co-immunoprecipitation experiments (Matakatsu et al., 2017; Zhang et al., 2016; Misra and Irvine, 2016). We further found that while the Ft ICD can recruit Dachs in the absence of App and Dlish, deletion of either the M or C domain prevents recruitment of Dachs unless App and Dlish are present (Fig. 7F, G and Fig S4F-H). This result suggests that multiple Ft ICD domains could be involved in forming a complex between Ft and the core complex. Using endogenously tagged alleles and surface labeling protocols we found that in wing discs under normal conditions core complex components accumulate with Ds, as has been suggested previously, but do not accumulate with Ft. However, in the absence of Dco, which phosphorylates the Ft ICD and promotes its ability to degrade Dachs, core complex components do accumulate with Ft at cell contacts. Thus, our results suggest that Ft degrades Dachs by first recruiting the core complex in a similar manner to Ds.

### Functions of the Ds intracellular domain

Using CRISPR we have performed an extensive structure/function analysis of the Ds ICD. Removal of the Ds ICD has no observable effect on the abundance or localization of either Ds or Ft, but does dramatically affect the core complex. Consistent with the idea that the Ds ICD promotes core complex activity, *ds^ΔICD^*mutants display marked undergrowth. Interestingly both Dachs and App lost their normal punctate appearance in *ds^ΔICD^* clones, suggesting a key role for Ds in organizing these puncta and that they normally promote Dachs function in promoting growth.

Fine scale structure/function studies mapped the domains responsible for growth control and interaction with the core complex. Previous studies identified three motifs (CM1-3) in the ICD that are conserved between *Drosophila* Dachsous and its vertebrate orthologs Dachsous 1 and Dachsous 2 (Hulpiau and van Roy, 2009). Using the Echinoid S2 cell assay and CRISPR deletion mutants *in vivo*, we found that domain D is necessary for core complex recruitment. Additionally, deletion of domain D leads to proximo-distal patterning defects and undergrowth, similar to mutation of core complex components. Domain D contains the highly conserved CM2 motif (Hulpiau and van Roy, 2009). In zebrafish, tetratrico peptide repeat containing protein 28 (Ttc-28) was shown to interact with the CM2 motif in Dachsous-1b (Chen et al., 2018). Although its functions have not been studied, a Ttc-28 homologue (CG43163) exists in *Drosophila*. Interestingly, while other regions of the Ds ICD are not necessary for core complex recruitment, they also display growth defects, suggesting that these regions might interact with other proteins such as Riquiqui, Lowfat and Myo31DF that have been implicated in Ds-Ft signaling (Degoutin et al., 2013; Mao et al., 2009; González-Morales et al., 2015). Consistent with this idea, we found that deletion of other domains, such as domain A, complement the wing growth and leg patterning defects of domain D deletions.

### Concluding remarks

A striking feature of Ft, Ds and core complex localization that is emphasized by our imaging of endogenously tagged components is that all of these proteins predominantly accumulate in distinct puncta along the junctional cortex. Previous experiments have shown that the mobility of Ft and Ds in these puncta is dependent on their interaction with one another (Hale et al., 2015), suggesting that each punctum consists of Ds and Ft accumulated in trans on adjacent cells. We show in addition that all components of the core complex localize to these puncta in association with Ds. Furthermore, their formation is not an inherent property of the core complex but rather is driven by Ft and Ds because in *ds ft* mutant cells cortical Dachs, Dlish and App all appear much more uniformly distributed along the junctional cortex. It is still not known how these puncta form, but our observations that 1) App localization is essentially normal in *ft^ΔICD^* mutants while Dachs is uniformly distributed at the junctional cortex (Fig. 6D-E) and 2) App is severely mislocalized in *ds^ΔICD^* cells (Fig. 4K) suggests that App is recruited to puncta by Ds. A possible model consistent with our observations is that App lateral movement within the plasma membrane is restricted through interaction with the Ds ICD where it then promotes accumulation of Dachs and Dlish (Fig. 9). When Ds is bound to Ft in trans these interactions result in stable puncta that promote growth (‘ON’ state), while in the absence of Ft binding Ds promotes removal of the core complex from the junctional cortex (‘OFF’ state). The Ft ICD similarly recruits these proteins but their interaction is more transient due to Ft-mediated degradation of Dachs at junctional cortex (‘OFF’ state). Further studies will be required to test this model, but it provides a framework for understanding the molecular and functional interactions between Ds, Ft and the core complex in regulating organ growth.

**Figure 9.**
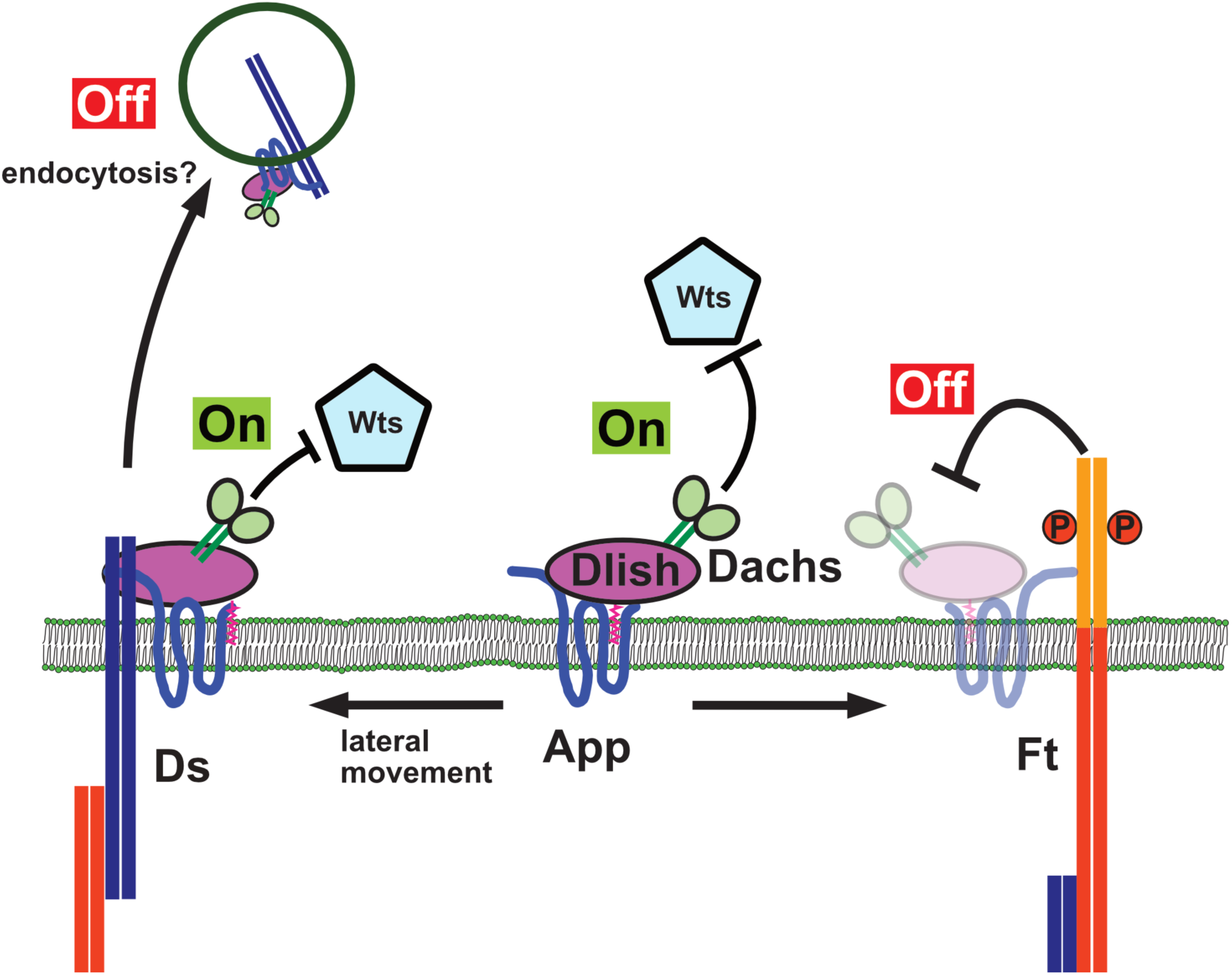
Regulation of junctional localization and abundance of the core complex by Ds and Ft. Dachs, Dlish and App interact at the junctional cortex to form the core complex and promote tissue growth by repressing the activity of Wts, the Hippo pathway kinase (’On’ state). When Ft is present, it binds the core complex and promotes degradation of Dachs (’Off’ state). Ds also binds the core complex, in particular App, and in the presence of Ft promotes the formation of stable core complex puncta that are protected from Ft-mediated degradation. However, in the absence of Ft, Ds facilitates removal of the core complex from the junctional cortex (possibly via endocyosis - see discussion) and in this way represses growth.

## Materials and methods

### Fly genetics

To induce mitotic clones, larvae were incubated at 37 ℃ for 40-120 min in a water bath (Xu and Rubin, 1993). The following stocks were used: *vasa-cas9* (Gratz et al., 2013; Sebo et al., 2014), *ds^UAO71^*(Adler et al., 1996), *ds^05142^*(BL11394)*, dachs^GC13^, dlish^Y003^* (Wang et al., 2019), *app^12-3^*, *app^DHHS^, ft^fd^ , ft^G-rv^, ft^sum^* (Bossuyt et al., 2014), *ft^61^* (Bosch et al., 2014), *ft^ΔICD^, ds^UAO71^ ft^G-rv^*, *ds^UAO71^ ft^fd^*, *hh-gal4, ban3-GFP, app^:YFP^, app^:V5^, dachs:GFP* (Bosveld et al., 2012), *ds:GFP* (Brittle., 2012), *ds^ΔICD:GFP^*, *ds^ΔICD^, ds^:Flag^, ds^ΔA:Flag^, ds^ΔΒ:Flag^, ds^ΔC:Flag^, ds^ΔD:Flag^, ft:GFP* (VDRC 318477), *V5:ft*, *HA:ds* (Ambegaonkar et al., 2012)*, UAS-dlish:Flag*, *UAS-dachs:V5* (Mao et al., 2006), *UAS-ds*, *UAS-app*, *ft RNAi* (VDRC 9396), *ds RNAi* (VDRC 36219), *app RNAi* (BL 35280), *y w hs-FLP; FRT 2A*, *y w hs-FLP*; *ubi-RFP FRT 2A*, *y w hs-FLP*; *ubi-GFP FRT 2A*, *y w hs-FLP; ubi-GFP FRT 40a*, *y w hs-FLP*; *ubi-RFP FRT 40a, TKO.GS00382* (BL79393) and *w^1118^*.

### Genotypes used for each figure

Fig. 1 H, I: *y w hs-FLP; ft^fd^ FRT 40a/ ds^UAO71^ ft^G-rv^ FRT 40a; ban3-GFP/ +*

Fig. 1 J: *y w hs-FLP; ft^fd^ FRT 40a/ ds^UAO71^ FRT 40a; dachs:GFP/ +*

Fig. 1 K: *y w hs-FLP; ft^G-rv^ FRT 40 a /ds^UAO71^ ft^fd^ FRT 40a; dachs:GFP/+*

Fig. 1 N: *y w hs-FLP; ft^fd^ FRT 40a/ ubi-mRFP FRT 40a; dachs:GFP*/ *dachs:GFP*

Fig. 2 D: *y w hs-FLP; ds^ΔICD:GFP^ FRT 40a/ ubi-GFP FRT 40a*

Fig. 3 C: *y w hs-FLP; ds^ΔICD^ FRT 40a/ ubi-mRFP FRT 40a; dachs:GFP/ dachs:GFP*

Fig. 3 F: *y w hs-FLP; ds^ΔICD^ ft^fd^ FRT 40a/ ubi-mRFP FRT40a; dachs:GFP/ dachs:GFP*

Fig. 4 E: *y w hs-FLP ft^fd^ FRT 40a/ ubi-mRFP FRT40a; app:YFP/ app:YFP*

Fig. 4 F: *y w hs-FLP; ds^UAO71^ FRT 40a/ ubi-mRFP FRT40a; app:YFP/ app:YFP*

Fig. 4 G: *w; ft RNAi/+; app:YFP hh-gal4/+*

Fig. 4 H: *w; ft RNAi/ +; app:YFP hh-gal4/ app RNAi*

Fig. 4 I: *w; ds RNAi* / +; *app:YFP hh-gal4/ +*

Fig. 4 J: *w; ds RNAi* / +; *app:YFP hh-gal4/ app RNAi*

Fig. 4 K: *y w hs-FLP; ds ^ΔICD^ FRT 40a/ ubi-mRFP FRT40a; app:YFP/ app:YFP*

Fig. 4 M: *w; app:YFP hh-gal4/ UAS-ds*

Fig. 5 X: *y w hs-FLP; ds^ΔD:Flag^ FRT 40a/ ubi-GFP FRT 40a; app:V5/ app:V5*

Fig. 6 D: *y w hs-FLP; ft^Δ^ ^ICD^ FRT 40a/ ubi-mRFP FRT 40a; dachs:GFP/ +*

Fig. 6 E: *y w hs-FLP; ft^Δ^ ^ICD^ FRT 40a/ ubi-mRFP FRT 40a; app:YFP/ app:YFP*

Fig. 6 F: *y w hs-FLP; ft^sum^ FRT 40a/ ubi-mRFP FRT 40a; app:YFP/ app:YFP*

Fig. 6 G: *y w hs-FLP; app:YFP FRT82 dco3/ app:YFP FRT82 ubi-RFP*

Fig. 6 H: *y w hs-FLP; HA:ds dachs:GFP FRT 2A/ ds+ FRT 2A*

Fig. 6 I: *y w hs-FLP; HA:ds app:V5 FRT 2A/ ds+ FRT 2A*

Fig. 6 J: *y w hs-FLP; V5:ft dachs:GFP FRT 2A/ ft+ FRT 2A*

Fig. 6 K: *y w hs-FLP; ft:GFP app:V5 FRT 2A/ ft+ FRT 2A*

Fig. 6 L: *y w hs-FLP; V5:ft dachs:GFP FRT 2A dco^3^/ ft+ FRT 2A dco^3^*

Fig. 6 M: *y w hs-FLP; ft:GFP app:V5 FRT 2A dco^3^/ ft+ FRT 2A dco^3^*

Fig. 8 A: *w; UAS-dachs:V5/+; hh-gal4 app:YFP/+*

Fig. 8 B: *w; UAS-app/+; dachs:GFP hh-gal4 /+*

Fig. 8 D: *y w hs-FLP; ds^UAO71^ ft^fd^ FRT 40a/ ubi-GFP FRT 40a*

Fig. 8 E: *y w hs-FLP; ds^UAO71^ ft^fd^ FRT 40a/ ubi-mRFP FRT 40a; app:YFP / app:YFP*

Fig. 8 F: *y w hs-FLP; ds^UAO71^ ft^fd^ dachs^GC13^ FRT 40a/ ubi-GFP FRT 40a*

Fig. 8 G: *y w hs-FLP; ds^UAO71^ ft^fd^ dachs^GC13^FRT 40a/ ubi-mRFP FRT 40a; app:YFP/ app:YFP*

Fig. 8 H: *y w hs-FLP; ds^UAO71^ ft^fd^ FRT 40a dlish^Y003^/ubi-GFP FRT40a dlish^Y003^*

Fig. 8 I: *y w hs-FLP; ds^UAO71^ ft^fd^ FRT 40a/ ubi-GFP FRT 40a; app^12-3^FRT 2A / app^12-3^ FRT 2A*

Fig. 8 J: *y w hs-FLP; ds^UAO71^ ft^fd^ FRT 40a/ ds^UAO71^ ft^G-rv^ FRT 40a; app^12-3^ FRT 2A/ ubi-GFP FRT 2A*

Fig. 8 K: *y w hs-FLP; ds^UAO71^ ft^fd^ FRT 40a/ ds^UAO71^ ft^G-rv^ FRT 40a; ubi-GFP FRT 2A / app^DHHS^ FRT 2A*

### Molecular cloning

To construct pMT-*ed:ds:Flag* and pMT-*ed:ft:Flag*, the pMT plasmid backbone and Echinoid extracellular domain was amplified from pMT-*ed:GFP:apkC* (Johnston et al., 2009) with iProof high-Fidelity DNA Polymerase (BIO-RAD). The intracellular and transmembrane domain of *ds* and *ft* were amplified from pUAST-*dsTMICD:Flag* (Zhang et al., 2016) or *Ft STI-4:FVH* (Feng and Irvine, 2009). DNA fragments were assembled with Gibson assembly. Deletions of these constructs were constructed using the Q5 site-directed mutagenesis kit according to the manufacturer protocol (NEB). For construction of pMT*-ed:Flag:vhh4*, the GFP nanobody sequence was amplified from pUAST-attB-NSlmb-vhhGFP4 (Caussinus et al., 2012, Addgene 35577) and fused to the Ed ECD as just described. All resulting plasmids were verified with DNA sequencing. Detailed information is available by request.

### Imaging

All imaging was done on fixed tissues or S2 cells. Fluorescently-tagged proteins were viewed using their native fluorescence. For other proteins, the following primary antibodies were used: Rat anti-Dachs (1:20,000; Matakatsu et al., 2017), Guinea pig anti-App (1:10,000; Matakatsu and Blair, 2008), Rabbit anti-Dlish (1:1,000; Zhang et al., 2016), Rabbit anti-Ds (1:1,000; Strutt and Strutt, 2002), Rat or Guinea pig anti-Ft (1:1,000; Sopko et al., 2009), Rat anti-*D*ECad (DCAD2, 1:1,000; DSHB; Oda et al, 1994) , Rat anti-Ci (2A1, 1:10; Motzny and Holmgren, 1995), Rabbit anti-HA (1:1,000; Abcam), Rabbit anti-HA (1:1,000; Y-11, Santa Cruz), iFluor 555 or iFluor 647 conjugated Mouse anti-V5 (1:100, GenScript), Mouse anti-Flag (1:20,000; M2, Millipore Sigma), Mouse anti-β-galactosidase (JIE7, 1:1,000; DSHB) and Alexa Fluor 647 Phalloidin (1:1,000; Thermo Fisher). Fluorescent secondary antibodies were obtained from Jackson laboratory or Invitrogen. After staining, wing discs were mounted in ProLong Diamond Antifade Mountant (Thermo Fisher). Images were taken using an LSM 880 confocal microscope (Zeiss, using Zen software), using Plan Apochromat 20×/0.8 na, Plan Apochromat 40×/1.4 na and Plan Apochromat 63×/1.4 na objectives. The acquired images were processed with Image J. Apical z-stacks were combined to make maximal projections using Image J.

App:V5 extracellular staining was performed as previously described (Hale et al., 2015). Wing discs were dissected at 5.5 h after puparium formation and incubated in iFluor 555 or iFluor 647 conjugated Mouse anti-V5 in dissection medium for 30 min on ice. After washing with dissection medium, wing discs were fixed with 2% formaldehyde in PBS for 10 min at room temperature, washed with PBS, and mounted. Images for whole wing discs were taken using tile scanning. Individual scanned images were aligned and combined into single images with Photoshop (Adobe) or Affinity designer (Serif).

To distinguish the native fluorescence of GFP and YFP tagged proteins in the same sample (Fig. 4A and C), we used the spectral detector on the Zeiss LSM 880 microscope to separate GFP from YFP signals. Additionally, we used the 458 nm line to excite GFP and the 514 line to excite YFP in separate tracks to minimize co-excitation of the fluorescent tags.

### In vitro studies

S2 cells were transfected according to Han (1996). pUAST constructs were co-transfected with pAWGal4 (kindly provided Y. Hiromi). To induce metallothionein promoter expression, CuSO_4_ was added to culture medium at a final concentration of 0.7 mM 24 h after transfection. S2 cells expressing pMT Ed constructs were allowed to aggregate as described previously (Matakatsu and Blair, 2004). After cells were agitated on a rotator at 70 rpm for 30-60 min, cells were attached on poly-L-lysine coated cover slips and fixed with 2 % formaldehyde in PBS for 10 min at room temperature. Cells were stained according to standard procedures.

### Immunoblotting and immunoprecipitation

Western blotting and immunoprecipitation are performed as previously described (Matakatsu et al., 2017). For immunoprecipitation of HA or Flag tagged proteins, anti-HA magnetic beads (Pierce) and anti-FLAG M2 agarose beads (Millipore Sigma) were used. Quantification of Dachs protein on immunoblots was performed as previously described (Matakatsu et al., 2017). The following primary antibodies were used for visualization of proteins: Rat anti-Dachs (1:10,000; Matakatsu et al., 2017), Mouse anti-αTubulin (1:20,000; Millipore Sigma), Guinea pig anti-App (1:10,000; Matakatsu and Blair, 2008), Mouse anti-Flag M2 (1:20,000; Millipore Sigma), Mouse anti-V5 (1:10,000; Genscript), Anti-V5 tag antibody [SV5-P-K](1:5,000; Abcam, ab206558) and Rabbit anti-GFP (1:2,000; kindly provided Dr. M. Glotzer). Blots were imaged using a LI-COR Biosciences Odyssey CLx. Efficiency of depletion with *app* and *dlish* dsRNA was confirmed by immunoblot (Fig. S4E). All experiments were performed at least three times and reproduced consistent results.

### Gene editing with the CRISPR-Cas9 system

The target sequence for CRISPR-Cas9 editing was designed according to flyCRISPR Optimal Target Finder (https://flycrispr.org/target-finder/). Target sequences were cloned into pU6-BbsI-chiRNA (Gratz et al., 2013; Addgene Plasmid 45946). The resulting plasmids were injected with or without donor plasmid into *vasa-Cas9* expressing embryos (Sebo et al., 2014, Gratz et al., 2013). Injections were performed by Genetivision.

To replace the Ds ICD with EGFP, EGFP was cloned between 5’ and 3’ homology arms into pBluescript II KS using Gibson assembly. The resulting plasmids and a pair of gRNA plasmids (pU6 target 4 for 5’ and pU6 target 1 for 3’) were co-injected. To obtain *ds* ICD deletions (Fig. 5A), the same pair of gRNA plasmids and appropriate donor plasmids containing Ds ICD deletions were co-injected.

To tag App with YFP, PCR amplified YFP sequence was cloned between 5’ and 3’ homology arms into pBluescript II KS (Fig. S1H). The resulting donor plasmid and gRNA plasmids (pU6 *app* target 2 and pU6 *app* target 3) were co-injected as before. The internal V5 tagging for extra cellular domain of App is summarized in Fig. S1I. gRNA plasmid (pU6 *app* V5 gRNA) and single stranded DNA oligonucleotides (*app* 3× V5) were co-injected as described. To make indel mutants for *ft^ΔICD^*, a gRNA plasmid (pU6 *ft* 5’ target 5) was injected as described (Fig. S4A). The resulting indel mutations caused premature stop codons after the transmembrane domain. After evaluation with anti-Ds staining, one mutant (24-3-5) was selected for further analysis.

After screening by genomic PCR with flanking primers, the isogenized lines were sequenced to determine the exact CRISPR event for all gene editing.

To generate *ds*^Δ*ICD*^ that lacks GFP fluorescence, the GFP coding sequence in *ds*^Δ*ICD:GFP*^ was disrupted by crossing with flies expressing *vasa-Cas9* and a *gRNA* for GFP (BL79393).

### Image Quantification

Wings from >20 females for each genotype were mounted on a slide glass in Permount (Fisher). Imaging was performed using a Zeiss Axioplan 2 microscope, using an EC Plan-Neo 5× objective. The images were taken using EOS REBEL T2i camera. The areas of wing blade were manually measured using ImageJ.

To quantify expression levels for App:YFP, Dachs:GFP and anti-Dlish staining in Fig. 8C, the mean gray value for a 25 x 25 µm square in the anterior (wild type) and posterior (overexpression) compartments in the proximal area of the wing disc (N>11) was determined using ImageJ.

### Statistical analysis

Minimum and Maximum are shown as bars in all box plots. All data points for box plots are shown as dots except Fig. 5M. Graphs were generated using GraphPad Prism 10. Statistical significances were evaluated using unpaired two-tailed t tests. Line scans were performed with ImageJ and quantified with Graph Pad Prism10.

### Online supplemental material

Fig. S1 shows anti-Ft staining for the wing disc displayed in Fig. 1J, anti-Dachs staining in *ds ft* and *ft* mutants, quantification of junctional intensity for Dachs in *ft* and in *ds ft* mutant, Ds distribution in *ft* clones shown in Fig. 4E, expression and subcellular localization of Ds:GFP and DsΔICD:GFP, tagging of endogenous App with CRISPR-Cas9 and co-localization of core complex proteins. Fig. S2 shows the amino acid sequence of the D domain in Ds and Dchs1 and the requirement of the D domain for core complex recruitment. Fig. S3 shows Ds staining in *ds* deletion mutants, colocalization of Ds deletions with Dachs, wing and leg phenotypes in trans-heterozygotes expressing different *ds* deletions. Fig. S4 shows the structure of the CRISPR-Cas9 induced *ft^ΔICD^* mutation, Dachs and App levels in the *ft^61^* mutant, phosphorylation of Ft and Ft^sum^ in S2 cells. Fig. S5 shows effects of *dlish* or *app* overexpression on core complex proteins, suppression of overgrowth loss of *app* or *dlish* in *ds ft* mutants, recruitment of Dachs by GFP tagged Dlish or App in S2 cells.

## Acknowledgements

We thank Drs. S. Blair, H. McNeill, D. Strutt, K. Irvine, Y. Bellaïche, G. Halder, I. Hariharan, J. Price, the Bloomington and Vienna Stock Centers, and the Developmental Studies Hybridoma Bank for reagents. We also thank M. Wang, W. J. Desrosiers for help with some experiments, and S. Blair, S. Tokamov and A. Smith for critical reading of the manuscript.

This research was funded by NIH research grant GM134128 to R. Fehon.

The authors declare no competing financial interests.

Author contribution: Conceptualization and methodology, H. Matakatsu and R. G. Fehon; Investigation, H. Matakatsu; Interpretation of data, H. Matakatsu and R. G. Fehon; Original draft, H. Matakatsu and R. G. Fehon; Writing, review and editing, H. Matakatsu and R. G. Fehon. Funding acquisition, R. G. Fehon, Project administration and Supervision, R. G. Fehon.

---

**Fig. S1.**
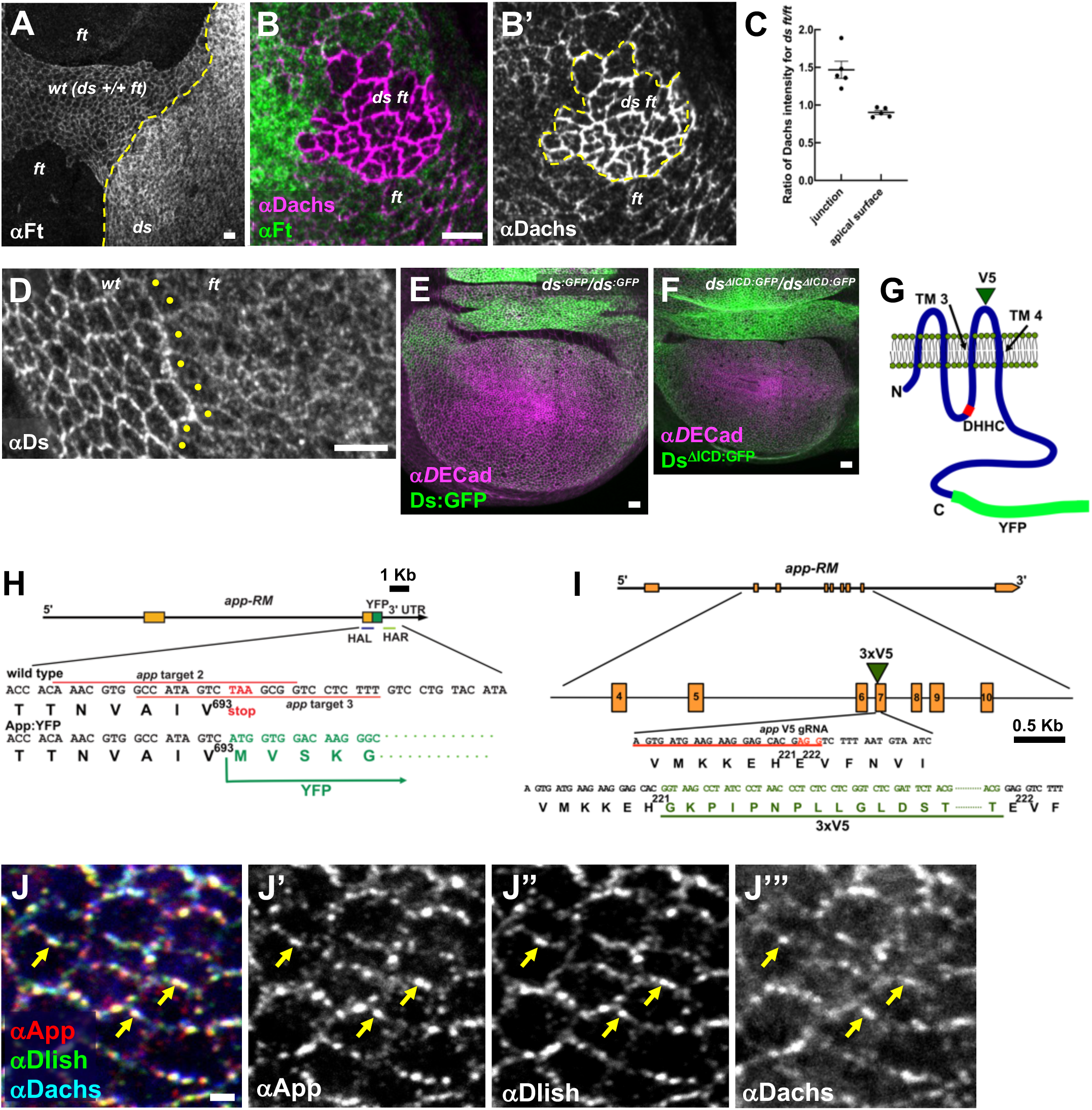
(A) anti-Ft staining for the wing disc displayed in Fig. 1J. *ft* clones are indicated by the lack of staining. The *ds* clone displays stronger Ft staining than wild type (*ds +/+ ft trans-*heterozygous) cells. (B) Comparison of Dachs subcellular localization between *ft* cells and *ds ft* cells (see also Fig. 1K-K’. The wing disc was stained with anti-Dachs staining). (C) Ratio of Dachs:GFP in *ds ft* compared to *ft* at junctions or the medial cortex. (D) anti-Ds staining for the wing disc displayed in Fig. 4 E. Ds is mislocalized at apical surface and junction. Expression and localization of Ds:GFP in *ds^:GFP^* homozygote (E) and DsΔICD:GFP in *ds^ΔICD:GFP^* homozygote (F) in wing discs. Adherens junctions are marked by anti-*D*ECad staining. Wing discs in *ds^ΔICD:GFP^* homozygotes are smaller than *ds^:GFP^* homozygotes. (G) App tagging by CRISPR-Cas9 gene editing. YFP was tagged at C-terminus. 3×V5 was inserted at extra cellular domain between transmembrane domains 3 (TM 3) and 4 (TM 4). (H) C-terminus tagging with YFP for *app* using CRISPR-Cas9. Target sequence (red line) and donor DNA are shown. HAL (blue line), left homology arm; HAR (light green line), right homology arm. (I) Internal V5 tagging for *app* using CRISPR-Cas9. 3×V5 (green triangle), target sequence (red underline), and PAM site (red letters) are shown. 3×V5 epitope tag was inserted after Histidine at 221. (J) Colocalization of App, Dlish, and Dachs at junctions. App, Dlish, and Dachs are colocalized at junctional puncta (yellow arrows). Scale bars, 5 µm (A, B, D), 10 µm (E, F), 1 µm (I).

**Fig. S2.**
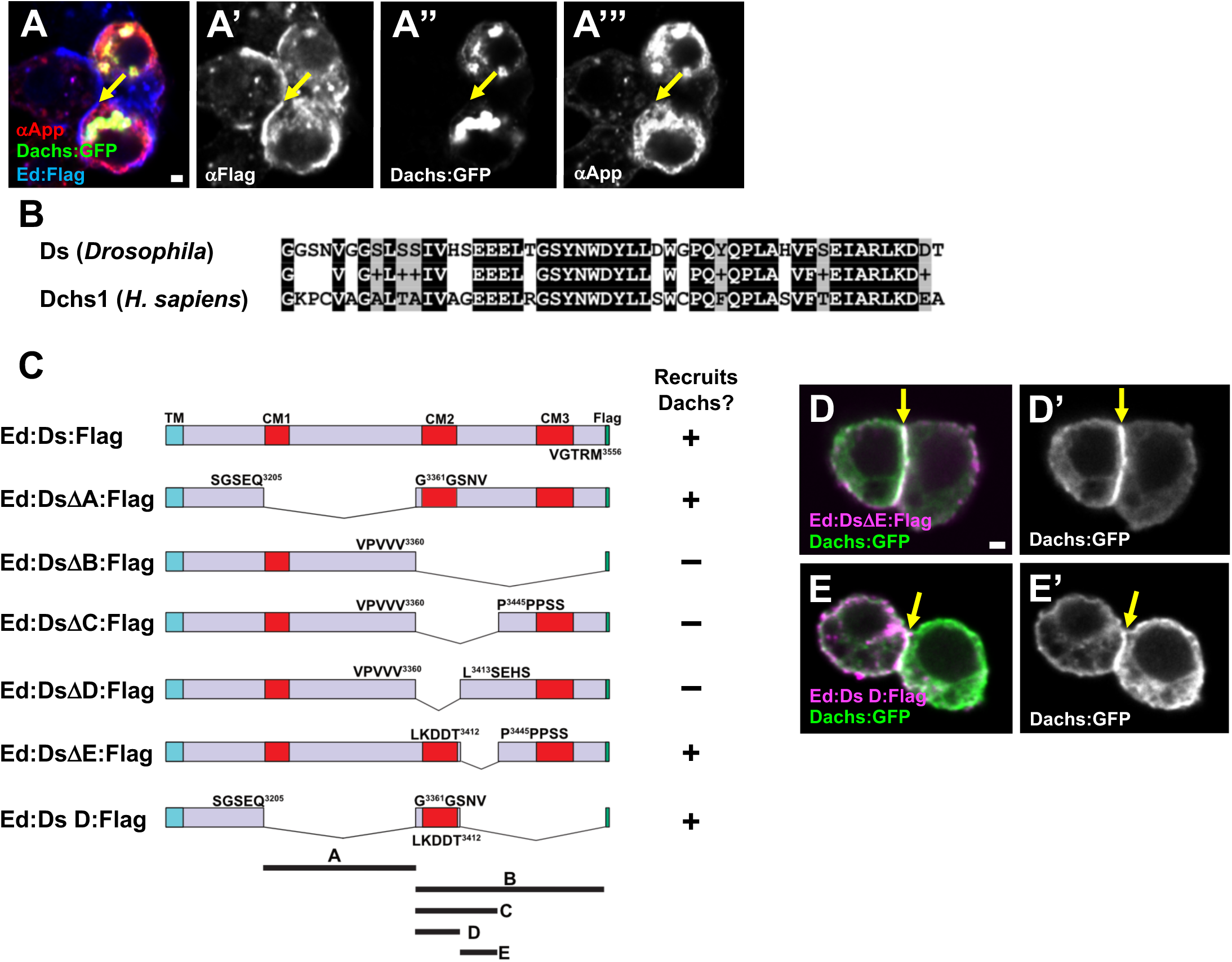
(A) Ed:Flag does not recruit Dachs:GFP and App to cell-cell contacts in S2 cells. Ed:Flag is localized at cell-cell contacts (yellow arrow inn A’). Dachs:GFP and App are distributed to cytoplasm and cytoplasmic foci (A’’, A’’’). (B) High homology between Ds and Dachs1 (*H. sapiens*) in domain D. Identical (black) and similar (gray) amino acids are shaded. (C) Constructs for Ed extracellular domain fused to Ds ICD or its deletions for the S2 cell recruitment assay. (D) Ed:DsΔE:Flag recruits Dachs:GFP to cell-cell contacts. (E) Ed:Ds D:Flag recruits Dachs:GFP to cell-cell contacts. Scale bars, 1 µm (A, D).

**Fig. S3.**
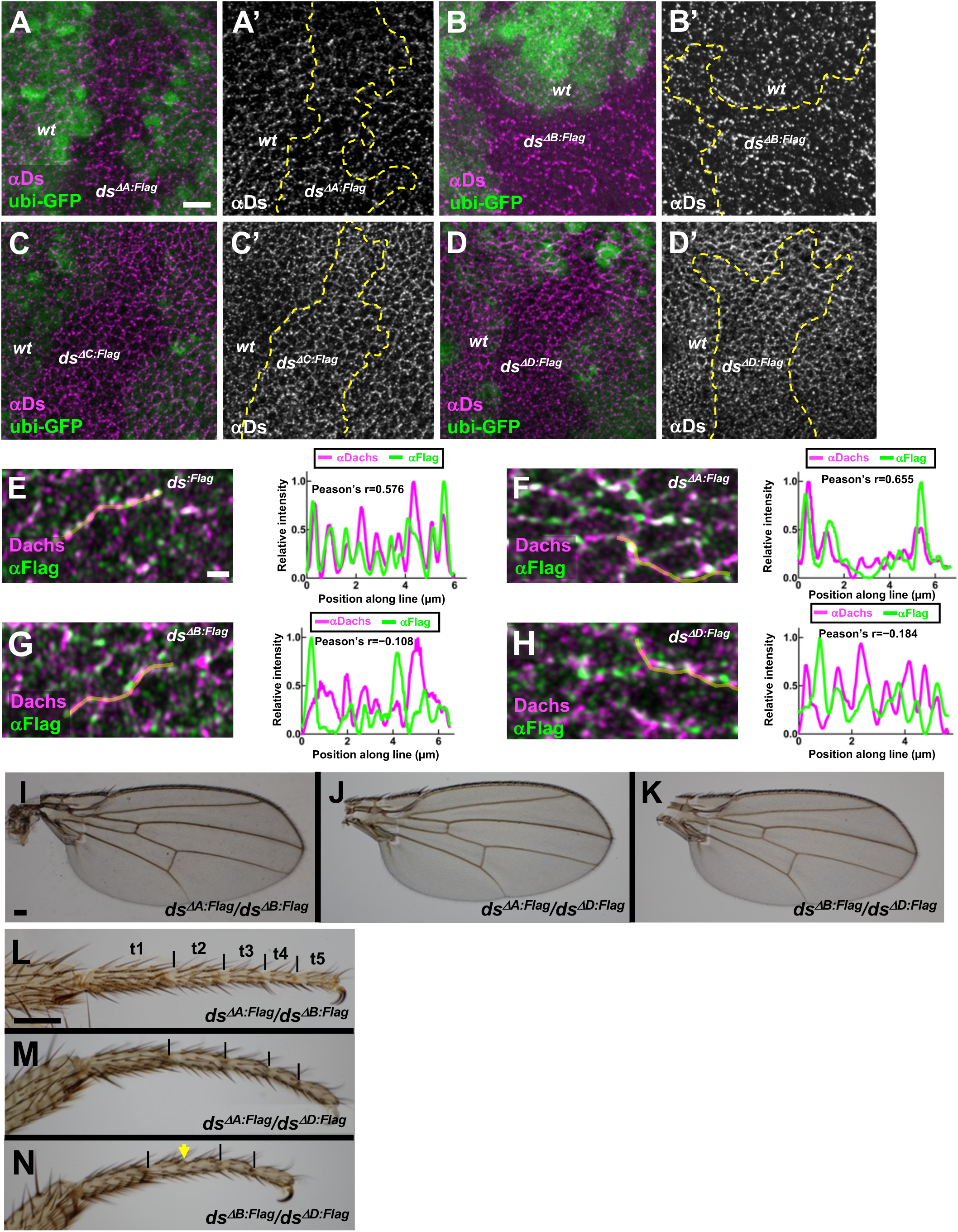
(A-D) anti-Ds staining in *ds^ΔA:Flag^*, *ds^ΔB:Flag^*, *ds^ΔC:Flag^* and *ds^ΔD:Flag^* clones. Mutant clones are labeled by the absence of GFP. The level and distribution of Ds did not change in *ds* deletion mutants. (E-H) Quantification of fluorescence intensity along cell junctions showing colocalization of Dachs with Ds:Flag and its deletions. Dachs:Flag (E) and DsΔA:Flag (F) show stronger colocalization with Dachs. DsΔB:Flag (G) and DsΔD:Flag (H) show weaker colocalization with Dachs. (I-K) Wings of the indicated genotypes showing transheterozygous complementation. *dsΔA:Flag/ dsΔB:Flag* and *dsΔA:Flag/ dsΔD:Flag* wings are almost normal size (I, J). *dsΔB:Flag/ dsΔD:Flag* wings display slight undergrowth (K). (L-N) Leg phenotype in transheterozygous combinations. Boundaries of tarsal segments are marked with black lines. *dsΔA:Flag/ dsΔB:Flag* (L) and *dsΔA:Flag/ dsΔD:Flag* (M) legs display normal segmentation. *dsΔB:Flag/ dsΔD:Flag* legs (N) are fused at T2 and T3 (yellow arrow). Scale bars, 5 µm (A), 1 µm (E), 100 µm (I, L).

**Fig. S4.**
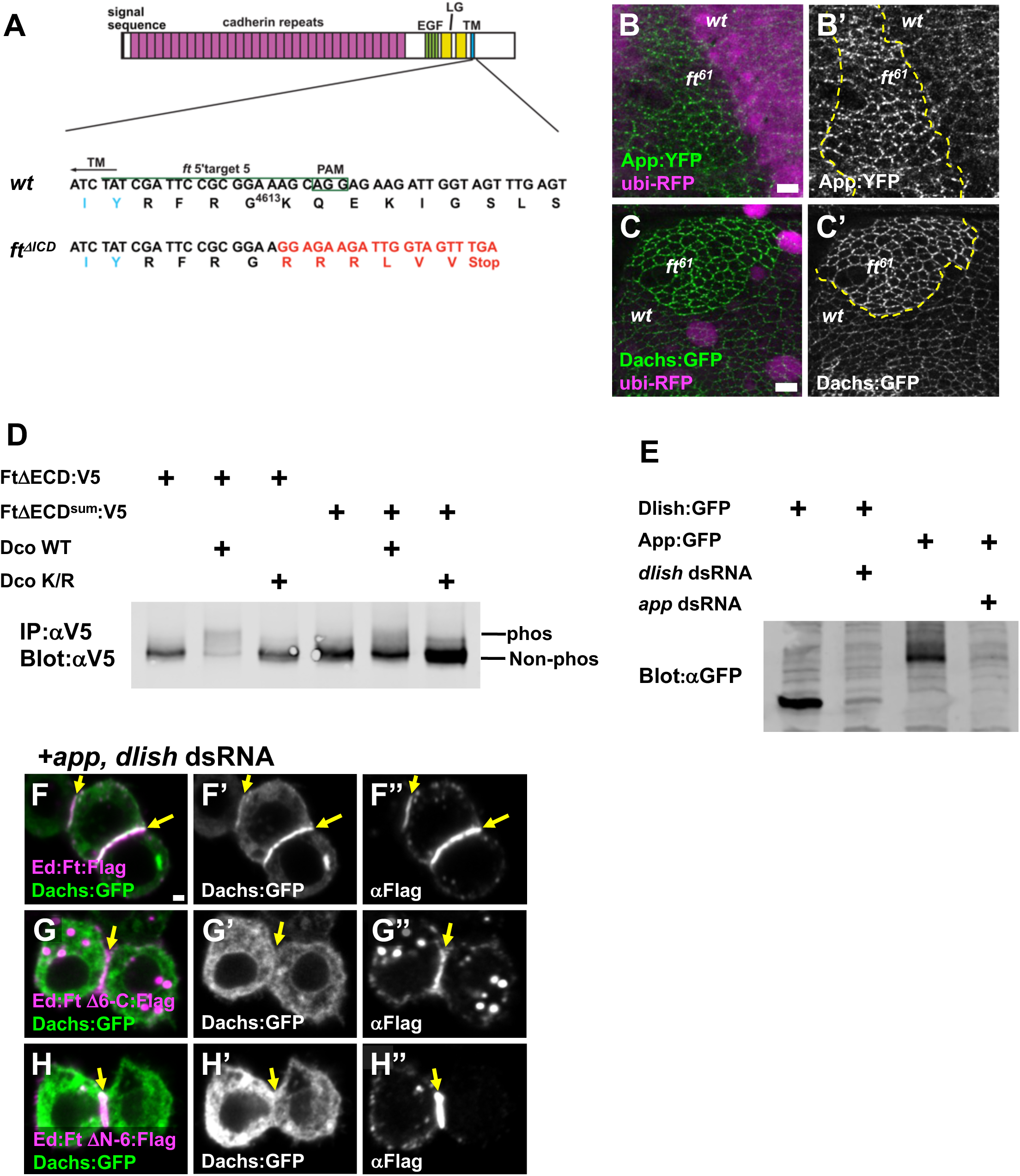
(A) CRISPR-Cas9 induced *ft^ΔICD^* mutant. The target sequence (green line), PAM site (green box) and transmembrane domain (blue letters) are shown. The resulting 4 bp deletion led to a premature stop codon after transmembrane domain. The altered amino acids are shown in red letters. (B) App:YFP is increased in junctional puncta in *ft^61^* mutant cells. (C) Dachs:GFP is increased at the junctional puncta in *ft^61^* mutant clones. Mutant clones are marked by absence of ubi-RFP. (D) Phosphorylation of wild-type Ft and Ft^sum^ by Dco in S2 cells. Dco WT promotes phosphorylation of Ft. The Ft^sum^ is less phosphorylated by Dco. (E) Efficacy of *app* and *dlish* dsRNAs in S2 cells. Both proteins are strongly reduced by co-transfection of dsRNA. (F) Ed:Ft:Flag recruits Dachs:GFP to cell-cell contacts in *app* and *dlish* depleted cells. (G) Ed:FtΔ6-C:Flag fails to recruit Dachs:GFP in *app* and *dlish* depleted cells. (H) Ed: FtΔN-6:Flag fails to recruit Dachs:GFP in *app* and *dlish* depleted cells. Scale bar, 5 µm (B, C) 1 µm (F).

**Fig. S5.**
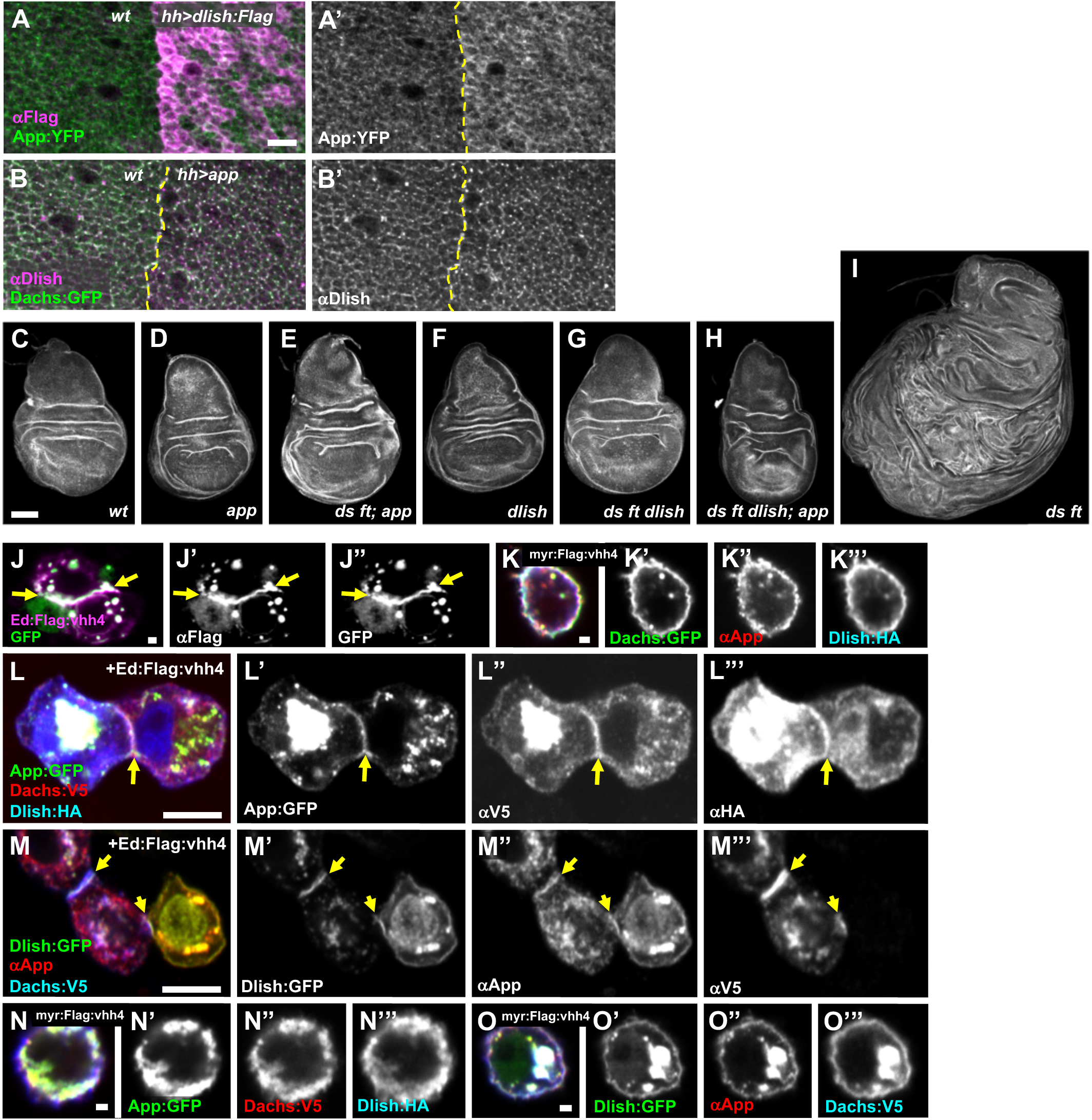
(A) App:YFP levels in *hh>dlish:Flag* at the junctional cortex. Dlish:Flag was expressed in the posterior compartment. App:YFP is increased with Dlish overexpression. (B) Dlish and Dachs:GFP in *hh>app.* Dlish and Dachs:GFP are enriched at junctional puncta with App overexpression. (C-I) Representative wing discs stained with phalloidin showing growth defects of the following genotypes: wild type (C), *app^12-3^* (D), *ds^UAO71^ ft^fd^; app^12-3^/ds^UAO71^ ft^G-rv^; app^12-3^* (E), *dlish^Y003^* (F), *ds^UAO71^ ft^fd^ _dlish_Y003_/ ds_UAO71 _ft_G-rv _dlish_Y003* _(G), *ds*_*UAO71 _ft_fd _dlish_Y003_; app_12-3_/ ds_UAO71 _ft_G-rv _dlish_Y003_; app_12-3* _(H) and *ds*_*UAO71 _ft_fd_/ ds_UAO71 _ft_G-^rv^* (I). Overgrowth in *ds ft* is partly suppressed with *app* or *dlish* mutant. (J) Recruitment of cytoplasmic GFP by Ed:Flag:vhh4. GFP is recruited to cell-cell contacts (yellow arrows). (K) Membrane tethering GFP nanobody recruits Dachs:GFP, App and Dlish to cell cortex. (L) Ed:Flag:vhh4 recruits App:GFP, Dachs and Dlish to cell-cell contacts (yellow arrow). (M) Ed:Flag:vhh4 recruits Dlish:GFP, App and Dachs to cell-cell contacts (yellow arrows). (N) Membrane tethering GFP nanobody recruits App:GFP, Dachs:V5 as well as Dlish:HA to cell cortex. (O) myr:Flag:vhh4 recruits Dlish:GFP, App and Dachs:V5 to cell cortex. Scale bars, 5 µm (A, L, M), 100 µm (C), 1 µm (J, K, N, O)

